# Cellular mechanisms underlying beneficial versus detrimental effects of bacterial antitumor immunotherapy

**DOI:** 10.1101/2024.02.15.580555

**Authors:** Jesse Garcia Castillo, Sebastian Fernandez, Timothy Campbell, Diego Gonzalez-Ventura, Jacob Williams, Julia Ybarra, Nicole Flores Hernandez, Elina Wells, Daniel A. Portnoy, Michel DuPage

## Abstract

Bacteria engineered to express tumor antigens as a cancer vaccine have yielded mixed results. Here, we utilized an attenuated strain of *Listeria monocytogenes* (*ΔactA, Lm*) that does not express tumor antigen to explore the immunological response to *Listeria* itself in the context of intravenous (IV), intratumoral (IT), or a combination of IV+IT administration into tumor-bearing mice. Unexpectedly, we found that *Lm* persisted in tumors of immune competent mice, regardless of the administration route. While IT *Lm* alone led to the recruitment of immunosuppressive immune cells that promoted tumor growth, IV *Lm* followed by IT *Lm* controlled tumor growth. IV *Lm* vaccination generated a pool of anti-*Lm* cytotoxic CD8 T cells that killed *Lm*-infected non-tumor cells to control tumor growth both indirectly, by limiting cancer cell proliferation, and directly, by enhancing tumor-specific T cell responses. Our findings reveal a differential impact of IT *Lm* administration on tumor progression that depends on the presence of anti-*Lm* CD8 T cells, which alone are sufficient to promote therapeutic efficacy.

## Introduction

At the turn of the 20th century, William Coley was the first to demonstrate the efficacy and application of bacterial cancer immunotherapies through combinations of intratumoral and systemic injections of cultured bacteria (gram-positive *Streptococci* and gram-negative *Serratia*) into patients with bone and soft-tissue sarcomas^1–3^. Today, local tumoral administration of Bacillus Calmette-Guerin (BCG), a live attenuated strain of the gram-positive intracellular pathogen *Mycobacterium bovis*, by intravesical injection is a standard of care for non-muscle-invasive bladder cancer and the only FDA approved remnant of bacterial cancer therapy^4^. Recently, the discovery that bacteria naturally colonize many solid tumors has uncovered the potential of bacteria to serve as a drug delivery system or even as a cancer detection probe^5–16^. Improved methods to genetically manipulate bacterial genomes has also made it feasible to engineer bacteria that express tumor antigens, cytokines or cytotoxic proteins to control cancer^17–21^. Hence, multiple breakthroughs in manipulating bacteria and their activity in the tumor microenvironment has reignited interest in the potential for effective bacterial-based cancer immunotherapies.

*Listeria monocytogenes* engineered to express tumor-associated antigens and injected intravenously has effectively controlled multiple cancer models in mice^22–26^. To date, *Listeria* engineered to express tumor antigens and injected intravenously is the most tested bacterial-based immunotherapy in clinical trials for cancer^27^. Despite strong evidence of the vaccine’s capacity to safely generate tumor antigen-specific CD8 T cells, the anti-cancer efficacy in patients has been mixed^28–33^. While several trials have shown improved outcomes in patients with otherwise untreatable cancers, such as malignant pleural mesothelioma or cervical cancer^28,30^, many *Listeria*-based vaccine trials have not shown significant improvement in tumor control compared to standards of care. Therefore, it is critically important to discern why these vaccines fail despite generating antitumor CD8 T cells. In contrast to other bacterial cancer therapies, whether *Listeria* directly colonizes tumors has not been adequately investigated, likely because colonization was never a goal for its use against cancer in patients. A better understanding of the dynamics of *Listeria* interaction with host immune cells, its migration, and its persistence in tissues may reveal vulnerabilities of the current *Listeria*-based strategies as well as lead to improved uses of *Listeria* as an immune adjuvant directly within tumors.

Effective antitumor immunity requires a robust pool of antitumor immune cells and a permissive tumor microenvironment (TME) that allows for antitumor immune cells to function^17,34–36^. However, the TME is often highly immunosuppressive, allowing cancer cells to evade the immune system despite the presence of tumor-specific cytotoxic CD8 T cells^37–40^. Suppressive immune cell populations such as regulatory T cells (Tregs) or myeloid derived suppressor cells (MDSCs) have been investigated as major obstacles that give rise to the immunosuppressive TME and counteract antitumor mechanisms by immunotherapies^4142^. Thus, the failure of *Listeria* vaccines might be due to *Listeria’s* singular use as a vaccine to generate antitumor CD8 T cells, without harnessing its capacity to stimulate the innate arm of the immune system and reverse immunosuppression directly within tumors. *Listeria* is rich in mechanisms to stimulate innate immunity directly in tumors^43^. Direct intratumoral injection of bacteria or bacterial products may have underlied the success of William Coley’s early therapies^44^. Recent preclinical studies as well as clinical trials have demonstrated the feasibility and efficacy of direct intratumoral injection of immune adjuvants naturally generated by *Listeria* and other bacterial species that can activate host cell pattern recognition receptors (PRRs) and stimulate the innate immune response against tumors^19,45–47^. However, there is a lack of understanding as to how these adjuvants or bacteria in the TME directly impact the tumor microenvironment by impacting different populations of immune cells.

Here, we show that either intratumoral (IT) or intravenous (IV) injection of *ΔactA*-attenuated *Listeria* (*Lm*) that is not engineered to express tumor antigens did not control tumor growth. Instead, IT *Lm* promoted tumor progression and *Lm* persisted predominantly in polymorphonuclear cells (PMNs) within the TME. *Lm* persistence in tumors was the result of IT *Lm* recruiting PMNs into the TME where they were polarized into a myeloid-derived suppressor cell phenotype (PMN-MDSC) that could suppress antitumor CD8 T cells^48,49^. In contrast, prior vaccination of mice against *Lm* by IV injection, either early in tumor development or prior to tumor initiation, led to strong tumor control upon IT *Lm* administration, even against highly aggressive sarcomas from genetically engineered mice. Tumor control required IFNγ and TNFα production as well as perforin-mediated killing by CD8 T cells specific to *Lm* antigen, which eliminated PMN-MDSCs harboring *Lm* within the TME. Killing of *Lm*-infected host cells mediated significant tumor control by reducing tumor cell proliferation and enhancing infiltration of antitumor CD8 T cells. In addition, the IV+IT dosing regimen increased the infiltration and reduced exhaustion phenotypes in tumor-specific CD8 T cells that further contributed to tumor control, especially at later time points after the IV+IT *Lm* regimen. Altogether, these results show that generating a localized anti-bacterial CD8 T cell response in tumors is sufficient to overcome the suppressive TME and inhibit tumor growth. Importantly, this work redefines the goals for effective anti-cancer bacterial therapies by revealing a critical role for direct T cell targeting of conserved bacterial antigens, which is more broadly applicable and rapidly translatable compared to engineering bacteria to express patient-tailored, tumor-specific neoantigens. These findings are fundamental to developing new and improved bacterial-based cancer therapies.

## Results

### Intratumoral injection of attenuated *Listeria* increases tumor growth

Immune adjuvants (i.e. PRR agonists) alone can be effective against cancers when delivered directly into tumors without the addition of engineered tumor antigens^45,50^ . To test whether antitumor responses could be generated by direct intratumoral (IT) injection of an attenuated strain of *L. monocytogenes*, which is not engineered to express tumor-specific antigens, we used an *!actA* mutant, denoted *Lm*, which contains the primary attenuating mutation used in many *Listeria* vaccine trials in patients^51^. *ΔactA Listeria* lack the ActA virulence factor, making the bacteria unable to polymerize host actin and spread from cell to cell, but importantly, *Lm* can still escape from phagosomes and grow in the host cell cytosol, making it a potent inducer of MHC-I-restricted CD8 T cell responses^52^. The *ΔactA* mutation renders *Lm* 1,000-10,000-fold less virulent than WT *L. monocytogenes* without decreasing its capacity to induce CD8 T cells. This feature has led to its use in many *Listeria*-based clinical trials due to its increased safety in patients^28,29,31,32^. However, the direct injection of *Listeria* into tumors of cancer patients has never been tested. We performed dose-escalating experiments with IT *Lm*, from 5×10^5^-5×10^7^ CFUs, into tumors 11 days post tumor inoculation, when tumors just became palpable (50-100mm^3^), and found no lethality at a dose of 5×10^7^ *Lm*. Therefore, we used 5×10^7^ CFUs for IT *Lm* injections in all experiments. IT injection of *Lm* did not result in tumor control in either MC38 or B16F10 tumor models but instead led to a significant increase in tumor growth **(Figures 1A-1C**, **Figures S1A-1B).** This result was unexpected, as *Lm* has been widely tested in experimental mouse models and was not observed to cause increased tumor growth. However, IT injection of *Lm* was never tested, so we hypothesized that this route of administration might be responsible for increased tumor growth. Therefore, we tested the impact of *Lm* administration on tumor growth after intravenous (IV) delivery, the most common method of administration, of 1×10^6^ *Lm* (a lower dose was required to prevent lethality by an IV route)^22^. In this scenario, we found no increase in tumor growth of B16F10 or MC38 tumors, nor did we observe tumor growth control using the *Lm* strain **(Figures S1C-1E)**. This finding is significant, as the route of *Lm* inoculation, IT versus IV, appears to convert *Lm* from either being detrimental or potentially therapeutic for the control of cancer.

**Figure 1.**
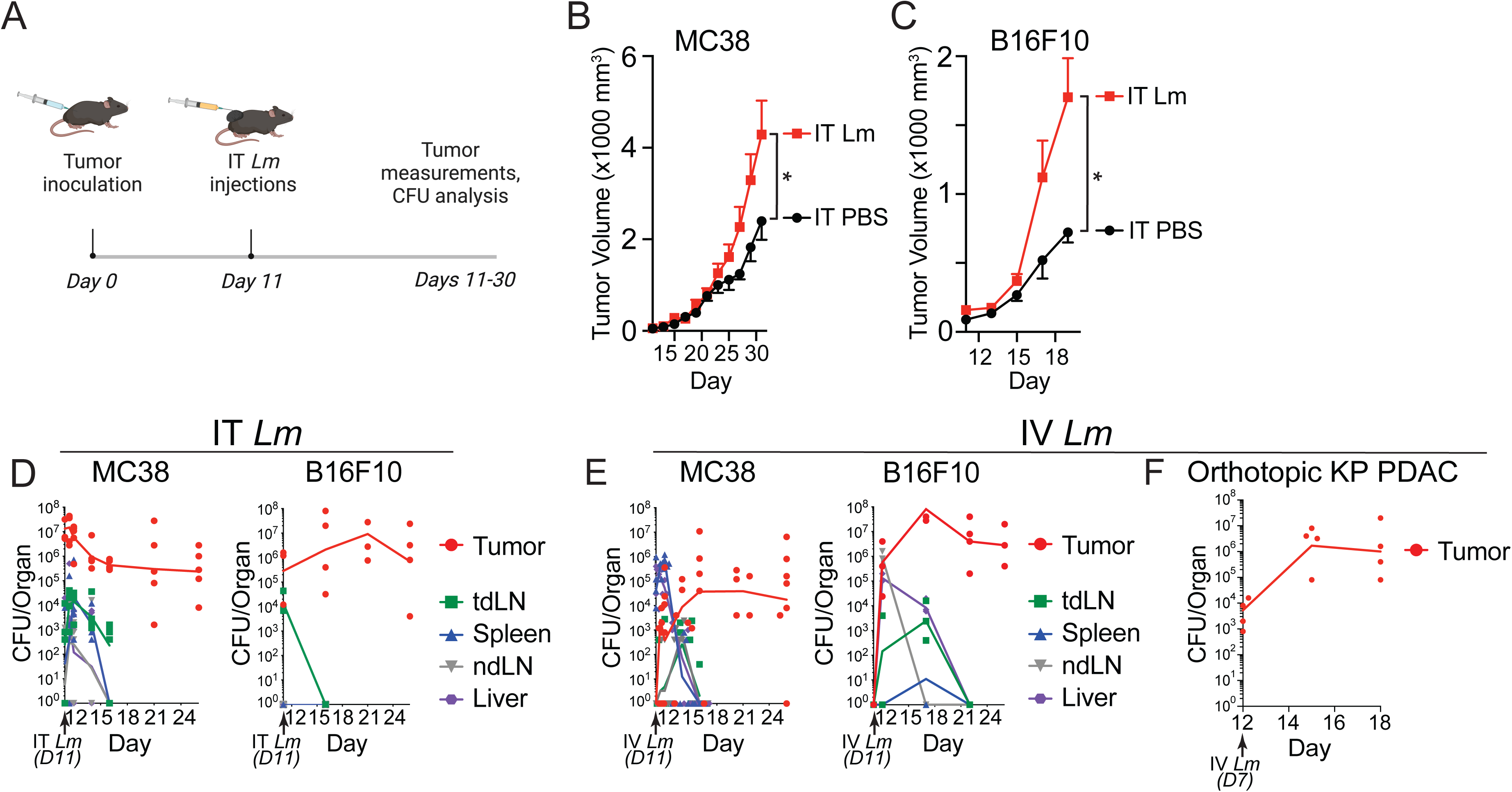
| Intratumoral injection of *Lm* leads to increased tumor growth and persistent colonization of *Lm* within tumors. (A) Experimental design. (B-C) C57BL/6J WT mice were injected with 5×10^5^ MC38 (B, data shown from n= 4-5 mice per group from one of three experiments) or 2×10^5^ B16F10 (C, data shown from n=4-5 mice per group from one of two experiments) tumor cells and then 5×10^7^ *Lm* CFUs IT 11 days later and tumors measured. (D-E) CFU analysis for *Lm* from indicated tissues after injection IT (D, 5×10^7^ CFU, n=4-8 mice pooled from two experiments) or IV (E, 1×10^6^ CFU, n=4-5 mice pooled from two experiments) of *Lm* 11 days after MC38 or B16F10 tumor inoculation. tdLN = tumor-draining lymph node, ndLN = non-tumor-draining LN. (F) CFU analysis for *Lm* at indicated time points from PDAC pancreatic tissue after orthotopic transplantation of FC1245^107^(n=4 mice). 10^6^ CFU of Lm injected on day 7. For all plots, *P<0.05, **P<0.01 by two-way ANOVA, mean ± s.e.m.

### *Lm* colonizes tumors and persists in immunocompetent mice

We hypothesized that the varying impacts on tumor growth after IT compared to IV *Lm* routes of administration could be due to varying *Lm* colonization in lymphoid organs or the tumor itself. To examine the localization of *Lm* in tumor-bearing mice after IV or IT *Lm* administration, we measured colony forming units (CFUs) for living bacteria from various tissues harvested from tumor-bearing mice at several time points after *Lm* administration. We found that *Lm* was cleared from spleens, lymph nodes, and liver by 5-10 days post injection in tumor-bearing mice, regardless of the route of delivery **(Figures 1D-E)**. However, *Lm* colonized subcutaneous MC38 and B16F10 tumors for up to 15 days, when experiments must end due to uncontrolled tumor growth (**Figures 1D-E**). *Lm* also colonized orthotopic pancreatic ductal adenocarcinomas (PDAC) after IV injection (**Figure 1F**)^53^. While other bacterial species have been shown to persist in tumors of mice, the persistence of live *Lm* within tumors of immunocompetent mice long-term (more than 3-5 days) has not been documented. The colonization and persistence of *Lm* in multiple tumor types after IV administration also revealed that systemically delivered *Lm* can seed, grow, and colonize tumors long-term, even if tumors are not easily accessible for direct IT injection.

### *Lm* infects PMN-MDSCs in tumors

We hypothesized that the distribution of *Lm* in specific cells of the tumor microenvironment after IT administration shaped the immune response to promote tumor growth. Because *Lm* is a facultative intracellular pathogen which primarily resides in host cells after infection, we sought to identify which cells harbor bacteria in the tumors of *Lm*-treated mice. To monitor intracellular infection of host cells by *Lm*, we used a fluorescent-protein based system wherein a Tag-RFP reporter was expressed under the control of the *actA* promoter in *ΔactA Listeria*, denoted *Lm-RFP* (**Figure 2A**)^54^. Since the *actA* promoter is induced more than 100-fold when *Lm* enters the host cell cytosol, expression of Tag-RFP indicates that the bacteria have entered the cytosol of a cell^55^. We observed co-localization of the RFP signal exclusively in CD45^+^ immune cells from single cell suspensions of MC38 or B16F10 tumors infected with *Lm-RFP*. Notably, we did not observe RFP fluorescence in CD45^-^ cells, which includes tumor cells (**Figure 2B**).

**Figure 2.**
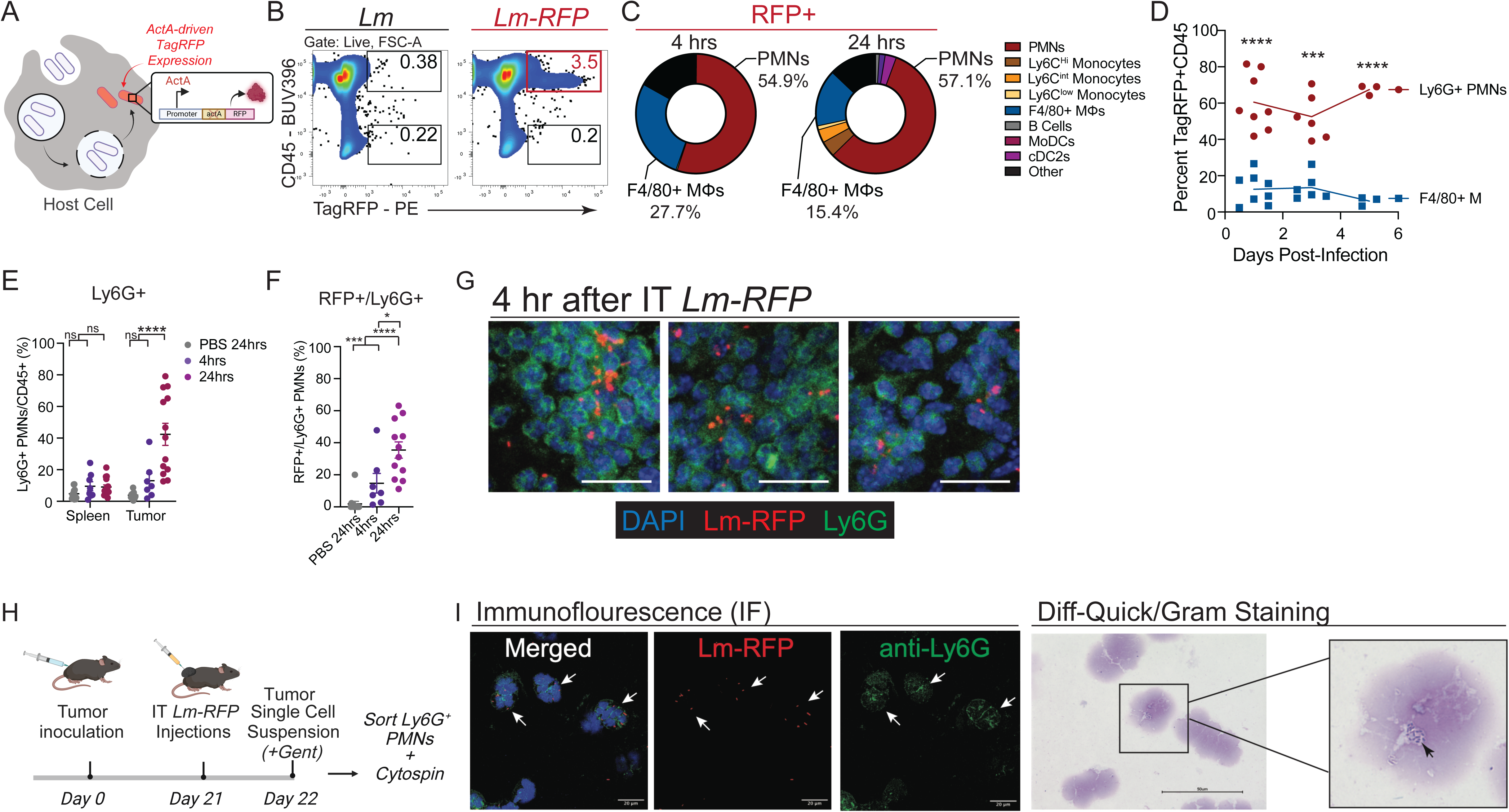
| *Lm* localizes in intratumoral PMNs. (A) Schematic of Tag-RFP expression from *Lm-RFP*. (B) Representative flow plots from single cell suspensions of MC38 tumors 4 hours post IT *Lm* (left) or IT *Lm*-*RFP* (right) showing CD45 staining versus Tag-RFP fluorescence. (C) Distribution of RFP^+^ immune cell types from MC38 tumors at 4 (left) or 24 (right) hours after IT *Lm* (representative data from three experiments with n= 3-6 mice/group). (D) Analysis of the fraction of PMNs and TAMs that were RFP^+^ at 1, 3, and 5 days after IT *Lm-RFP* in MC38 tumors (representative data from two experiments with n= 3-6 mice/group). (E) Frequency of Ly6G^+^ PMNs in spleen or tumors from MC38-bearing mice at 4 and 24 hours after IT PBS or IT *Lm*. (F) Frequency of intratumoral Ly6G^+^ cells that were RFP^+^ at 4 and 24 hours after IT PBS or IT *Lm*. Results in E-F are pooled from three experiments with n= 2-5 mice/group per experiment. (G) Images of MC38 tumors 4hrs after IT *Lm*: DAPI (blue), *Lm-RFP* (red), Ly6G (green). Scale bar = 20 μm. (H) Schematic for cytospin imaging analysis of sorted intratumoral Ly6G+ PMNs 24hrs after IT *Lm*. (I) Immunofluorescence (right) and Diff-Quick gram staining of sorted intratumoral Ly6G+ PMNs 24hrs after IT *Lm*. For immunofluorescence: DAPI(blue), Lm-RFP (red), Ly6G (green). Scale bar = 20 μm. For Diff-Quick: Scale bar = 50 μm For all plots, mean ± s.e.m. and *P<0.05, **P<0.01, ***P<0.001 by two-way ANOVA (D-E) or student t-test (F, G, H).

Next, we tracked which types of CD45^+^ immune cells harbored *Lm* in their cytosol. At 4-and 24-hours post-infection, we observed an enrichment of RFP-expressing bacteria in phagocytic immune cells, especially Ly6G^+^ polymorphonuclear (PMN) cells in both MC38 and B16F10 tumors models (**Figures 2C and S2A-D**). To a lesser extent, *Lm-RFP* was detected in F4/80^+^ macrophages, Ly6C^+^ monocytes, and CD11c^+^MHC-II^+^ dendritic cells (**Figures 2C and S2B**). This distribution of *Lm* in phagocytic immune cells, primarily PMNs and macrophages, was maintained after the initial infection for up to five days (**Figure 2D**). However, in comparison to tumors at steady state, IT *Lm* only induced a significant influx of PMNs into tumors, whereas other phagocytic cell frequencies were unchanged (**Figures 2E and S2C-E)**. The RFP^+^ PMNs also increased between 4 and 24 hours after IT delivery (**Figure 2F and S2F**).

To rule out that *Lm* was phagocytosed during the single cell preparation for flow cytometry analysis, we performed several experiments. First, we used confocal microscopy of frozen tumor sections and also observed cytosolic RFP+ *Lm* within Ly6G+ PMNs that aggregated in clusters within tumors (**Figure 2G and Figure S2G**). Second, we prepared single cells from tumors in the presence of the antibiotic gentamicin - to kill extracellular *Lm*, FACS purified Ly6G^+^ PMNs, and performed microscopic analysis of cytospun cells **(Figure 2H)**. We observed cytosolic RFP+ *Lm* in Ly6G^+^ PMNs by fluorescence microscopy with anti-Ly6G and by separate Gram staining **(Figure 2I)**. Altogether, these results demonstrate that PMNs serve as a reservoir for *Lm* in tumors after IT injection. These results were unanticipated, as we expected PMNs would rapidly kill phagocytosed *Lm*, preventing their escape into the cytosol and the expression of RFP^56–58^.

### IT *Lm* recruits PMNs that differentiate into pro-tumorigenic PMN-MDSCs within the tumor microenvironment

We hypothesized that the colonization and persistence of *Lm* in the cytosol of PMNs was the result of PMN differentiation into myeloid derived suppressor cells (MDSCs) within tumors, which could also promote tumor progression by inhibiting T cell responses^32,59–61^. Phenotyping of PMNs at 4 and 24 hours after IT *Lm* revealed a dynamic increase in the frequency of PMNs expressing the prototypical PMN-MDSC marker CD14, and CD14 positivity was enriched in RFP^+^ cells (**Figures 3A-B and S3A-B**)^62^. To further confirm differentiation of PMNs into MDSCs within tumors, we sorted Ly6G^+^ PMNs from MC38 tumors and analyzed the expression of additional genes known to be upregulated in MDSCs by qPCR. We observed that Ly6G^+^ cells sorted from tumors had higher expression of *Cd274*, *Cd14*, and *Vegfa* transcripts in comparison to their counterparts sorted from the spleen or bone marrow (**Figure 3C**). These results suggest that *Lm* infection within tumors recruits PMNs that then differentiate into an MDSC phenotype in tumors, which may make them a permissive reservoir for *Lm* colonization.

**Figure 3.**
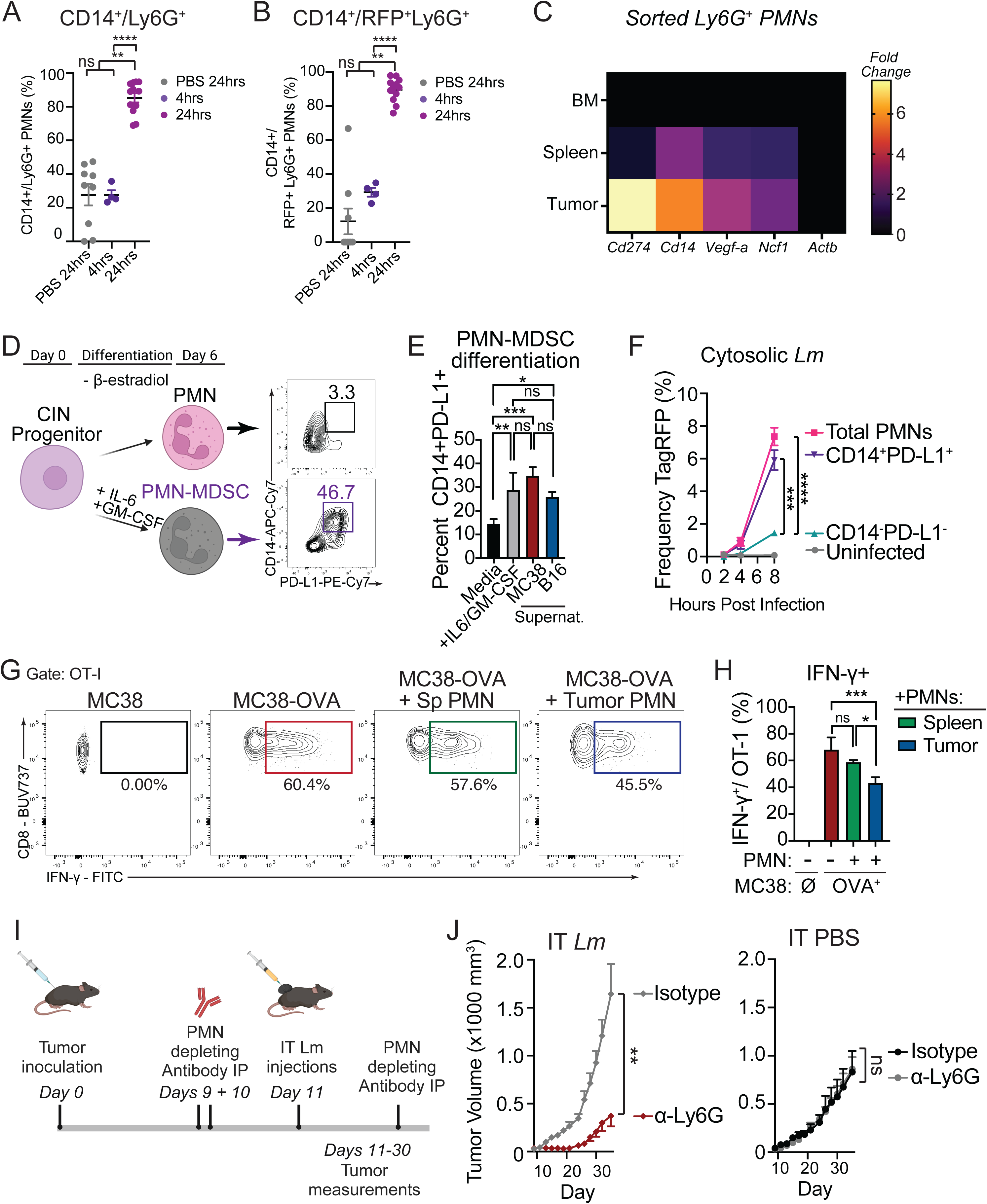
| PMN conversion into MDSCs permits *Lm* infection and promotes tumor growth. (A-B) Frequency of CD14^+^ cells amongst Ly6G^+^ (A) or RFP^+^Ly6G^+^ cells (B) from MC38 tumors 4 and 24 hours after IT PBS or IT *Lm*. Results are pooled from three experiments with n= 2-5 mice/group per experiment. (C) Expression of indicated genes by qPCR analysis from bone marrow (BM), spleen, or tumor of MC38-bearing mice. Results from n= 6 mice from a single experiment. (D) Experimental strategy and representative flow plots for *in vitro* differentiation of conditionally immortalized neutrophil (CIN) progenitors into PMNs (top) or PMN-MDSCs (bottom). (E) PMNs derived from CINs were cultured with conditioned supernatants from 3T3, MC38, or B16F10 cells for 24 hours *in vitro* and the frequency of CD14^+^PD-L1^+^ cells was assessed compared to unconditioned media or IL6+GM-CSF treatment (representative of two experiments). (F) Heterogeneous CD14^+/-^ PMNs differentiated from CINs as in (A) were infected with *Lm-RFP in vitro* and Tag-RFP^+^ cells were identified 2, 4, or 8 hours later in all cells (Total PMNs), CD14^+^ cells, or CD14^-^ cells by flow cytometry (representative of three experiments). (G) Representative flow plots of IFNγ production from OT-I CD8 T cells after 24 hour co-culture with MC38 or MC38-OVA tumors in the presence or absence of FACS-purified PMNs from spleens or tumors of IT *Lm*-treated MC38-bearing mice. (H) Quantification of the frequency of IFNγ^+^ OT-1 CD8 T cells (G). Cells were plated at a 1:1 ratio of OT-1 T cells to PMNs. Results from 2 independent experiments. (I) Experimental design to deplete PMNs prior to IT PBS or IT *Lm* and measure tumors. (G) (J) Tumor growth measured in mice injected IT with *Lm* (left) or IT PBS (right) and either an isotype control antibody or anti-Ly6G-depleting antibodies prior to tumor inoculation and throughout the experiment (n=5 mice/group from one of two independent experiments). For all plots, *P<0.05, **P<0.01, ***P<0.001 by one-way Anova (B, F) or two-way ANOVA (C, E), mean ± s.e.m for *in vivo* experiments (E) or mean ± s.d. for *in vitro* experiments (B-C, G).

To test whether PMN-MDSCs become permissive to *Lm* growth in their cytosol compared to PMNs, we used conditionally immortalized neutrophils (CINs) or primary bone marrow progenitors differentiated into PMNs *in vitro* (**Figures 3D and S3C**)^61,63^. First, CIN-derived PMNs treated with conditioned supernatants from MC38 or B16F10 tumor cells increased expression of CD14 and PD-L1 relative to noncancerous, immortalized NIH 3T3 cells, indicative of a PMN-MDSC phenotype (**Figures 3D-E**). As a positive control, PMNs were differentiated into PMN-MDSCs by adding IL-6 and GM-CSF, which led to a similar induction of CD14 and PD-L1 in CINs and bone marrow derived PMNs (**Figures 3D** and **S3C**)^63^. To test whether the adoption of this MDSC phenotype increased susceptibility to *Lm* infection, we infected heterogeneous PMN cultures *in vitro* with *Lm-RFP* and monitored infection by flow cytometry. The CD14^+^ PMNs were almost exclusively associated with *Lm* cytosolic infection as determined by RFP expression (**Figures 3F and S3D**). These results strongly suggest that IT *Lm* recruits PMNs to tumors where they become converted into PMN-MDSCs that allow for *Lm* to infect and persist intracellularly in PMN-MDSCs.

PMN-MDSCs are widely associated with decreased antitumor immunity and increased tumor progression. MDSCs have been shown to promote tumor growth by inhibiting antitumor CD8 T cell responses^35,64^. To test whether PMN-MDSCs inhibited CD8 T cells to promote cancer progression, we co-cultured tumor-specific CD8^+^ T cells (*in vitro* activated OT-I T cells) with OVA-expressing MC38 tumor cells in the presence of PMNs sorted from spleens or MC38 tumors. The addition of Ly6G^+^ PMNs from tumors (PMN-MDSCs), but not spleens, led to a significant reduction in IFNγ cytokine production from OT-I T cells in response to direct recognition of OVA-expressing MC38 tumors (**Figure 3G-H**). Next, we tested whether the depletion of all PMNs prior to IT Lm infection could reverse enhanced tumor progression (**Figure 3I**). Indeed, depletion of Ly6G^+^ cells prevented increased tumor growth with IT *Lm* **(Figures 3J** and **Figure S3E-F).** Notably, anti-Ly6G had no effect on tumor growth if mice were not administered IT *Lm*. Interestingly, PMN depletion did not reduce bacterial colonization of tumors as measured by CFU analysis (**Figure S3G**), although, using *Lm-RFP*, we found *Lm* was now present primarily in macrophages (**Figure S3H**). These results support that IT *Lm* recruits PMNs to tumors where they are converted to a MDSC phenotype that promotes increased tumor growth. Therefore, *Lm* injection into tumors can promote tumor growth, warranting caution when using bacterial cancer therapies.

### A combination of IV+IT *Lm* leads to tumor control across multiple cancer models

As neither IV nor IT administration of *Lm* alone provided strong tumor control, we considered that both routes of infection together might generate therapeutic efficacy. We reasoned that IV injection of *Lm* would effectively generate adaptive T cell responses against *Lm*^65^, and that subsequent IT *Lm* injection would then recruit anti-*Lm* T cells into tumors to augment the antitumor immune response. To test this hypothesis, we inoculated mice with MC38 or B16F10 tumors and administered *Lm* IV four days later. At 11 days after tumor inoculation, when tumors were palpable, we injected *Lm* IT into tumors as done previously with IT *Lm* alone **(Figure 4A)**. This combination of IV *Lm* followed by IT *Lm* administration now significantly controlled both MC38 and B16F10 tumors **(Figures 4B-C and Figure S4A-B)**. Tumor control was evident by 9 days after IT *Lm* injection, and tumors remained controlled throughout the experiment. Overall, of the experiments performed, only IV+IT *Lm* administration led to tumor rejection (20% of mice, 8 out of 40) and tumor rejection was never observed with IV or IT *Lm* alone. Because IV administration of *Lm* also led to colonization of tumors (**Figure 1E)**, we tested whether a second IV administration of *Lm* could also control tumors without requiring direct IT *Lm* injection. We found that IV+IV also controlled tumors at two different doses of IV *Lm* (**Figure S4C).** These results indicate that a second dose of *Lm* IT or IV can control tumors after colonizing tumors without requiring the *Lm* strain to express tumor antigens. In addition, we found that while Ly6G^+^ PMNs were recruited equally to tumors one day after IT *Lm* administration in both the IV+IT and IT-only *Lm* dosing regimens, the frequency of Ly6G^+^ cells was significantly reduced by 5 days after IT *Lm* administration in the IV+IT setting **(Figure 4D)**. Furthermore, the fraction of CD14^+^ PMNs was almost completely abolished 5 days after IT *Lm* administration in the IV+IT *Lm* dosing regimen, a reduction even below their frequency in PBS-treated control tumors despite their robust increase 24 hours after IT *Lm* **(Figure 4E)**. Finally, when we depleted PMNs prior to IT *Lm* administration in the IV+IT *Lm* regimen, we did not observe any enhancement in tumor control, suggesting that the PMN reductions with IV+IT *Lm* are sufficient to prevent PMN-MDSC promotion of tumor growth after the IT *Lm* dose in the IV+IT therapeutic regimen **(Figure S4D)**.

**Figure 4.**
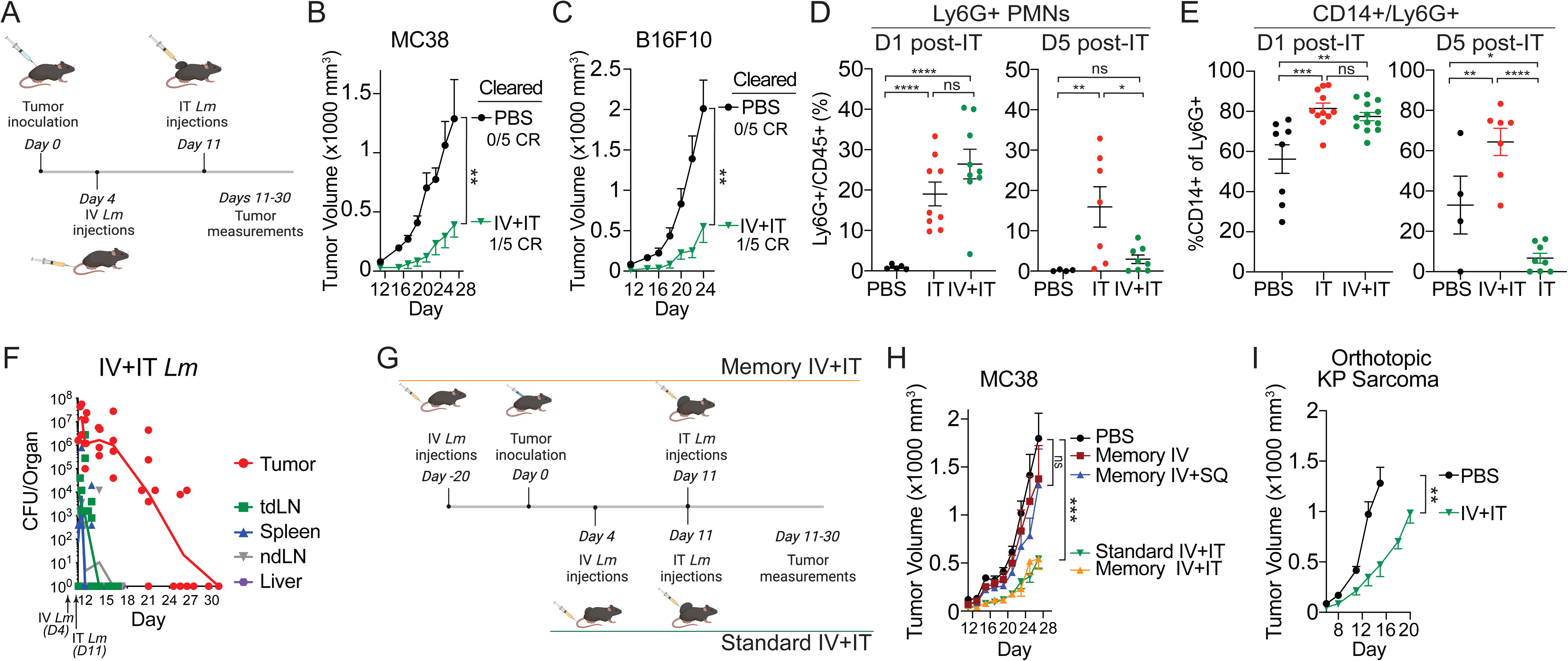
| IV+IT *Lm* controls multiple tumor types while clearing intratumoral bacteria and PMN-MDSCs from tumors. (A) Experimental strategy for IV+IT *Lm* dosing regimen. (B-C) IV+IT PBS versus IV+IT *Lm* regimen and growth of MC38 (B, data shown from n= 4-5 mice per group from one of three experiments) and B16F10 (C, data shown from n= 5-6 mice per group from one of two experiments) tumors. (D) Frequency of Ly6G^+^ PMNs of all CD45^+^ cells one day (left) and 5 days (right) after *Lm* IT in IV+IT *Lm* regimen (data shown from n= 5 mice per group from two pooled experiments). (E) Percent of Ly6G^+^ PMNs that were CD14^+^ 1 day (left) and 5 days (right) after IT *Lm* in IV+IT *Lm* regimen. (F) CFU analysis for *Lm* after IV+IT *Lm* regimen in MC38 tumor-bearing mice. Organs were harvested 4 hours, 12 hours, and on days 1, 3, 5, 10, 15, and 20 after IT *Lm* administration. (G) Experimental design for comparing prophylactic IV Lm (memory IV+IT) to standard IV+IT *Lm* dosing regimen. (H) MC38 tumor growth in the setting of memory IV+IT, memory IV only, and memory IV+SubQ *Lm* (injected on opposing flank) compared to PBS treatment or standard IV+IT *Lm*. (I) Orthotopic KP sarcoma growth with PBS versus a memory IV+IT *Lm* dosing regimen. For all plots, *P<0.05, **P<0.01, ***P<0.001 by student t-test (C, D, E) or two-way ANOVA (B, C, H, I), mean ± s.e.m.

Next, we wanted to establish whether *Lm* persisted in tumors in the setting of IV+IT *Lm* administration. While *Lm* persisted in approximately 80% of IV+IT *Lm*-treated tumors for up to 10 days after IT *Lm* administration, *Lm* was ultimately cleared from tumors, with CFUs dropping to below the level of detection between days 15-20 after IT *Lm* (day 21-30 of tumor growth) **(Figure 4F)**. We hypothesized that the clearance of *Lm* from tumors was the result of *Lm*-specific T cells that were generated by IV *Lm* and then recruited to tumors when *Lm* was injected IT. Therefore, tumor control with the IV+IT *Lm* dosing could be the result of the recruitment of *Lm*-specific T cells to tumors. Alternatively, because *Lm* colonized and persisted within the TME after IV administration at day 4 (**Figure S1F**), the early seeding of tumors might contribute to tumor control by changing the TME to augment the second IT injection of *Lm* in the IV+IT regimen. To address this possibility, we performed experiments in which IV *Lm* was administered prophylactically, 20 days prior to tumor inoculation (**Figure 4G**). In this setting, *Lm* was completely cleared from mice prior to tumor inoculation and therefore absent from tumors when IT *Lm* was administered. Now we could test whether recalled *Lm*-specific adaptive immune cells, rather than changes in the distribution or amount of *Lm* in tumors, was responsible for the antitumor efficacy of the IV+IT regimen. This experiment showed clearly that mice injected with *Lm* prophylactically (memory IV+IT) controlled tumors equivalently to the standard IV+IT regimen **(Figures 4H** and **S4E)**. However, prophylactic IV *Lm* injection alone, or IV *Lm* followed by a subcutaneous injection of *Lm* on the contralateral flank of mice (memory IV+SQ), did not control tumors **(Figure 4H)**. These results suggested that IT administration of *Lm* mediated antitumor immunity by recalling an adaptive immune response against *Lm* locally within the tumor.

Finally, we tested the IV+IT *Lm* dosing regimen in a clinically relevant mouse sarcoma model - the tumor type most often treated by Coley, and a clinically relevant model amenable to direct IT injections^1,23,66^. Using an aggressive *Kras^G12D/+^;p53^fl/fl^*-generated orthotopic sarcoma model^67^, we found that memory IV+IT *Lm* led to significant sarcoma control from *Lm*-treated sarcomas compared to tumors treated with PBS that had received IV *Lm* prophylactically **(Figures 4I and S4F)**. Together, these results showed that IV+IT *Lm* can clear *Lm* from tumors and control tumor growth in a manner that is not recapitulated with IV or IT *Lm* administrations alone.

### CD8 T cells are required for tumor control and *Lm* clearance with the IV+IT *Lm* **dosing regimen.**

The reduction of tumor growth that occurred during the IV+IT *Lm* treatment regimen compared to single-dose IV or IT *Lm* administrations suggested that an adaptive immune response mediated tumor control. We hypothesized that T cells were the critical adaptive immune cells important for tumor control in the IV+IT dosing regimen due to their established role in both anti-*Lm* and antitumor immunity^65,68,69^. Indeed, flow cytometric analysis of tumors 5-8 days after IT *Lm* injection (early) revealed an increase in the frequencies of both intratumoral CD8 and CD4 T cells in mice receiving IV+IT *Lm* compared to IV-only or IT-only *Lm* injections or PBS control injections (**Figures 5A-C**, **S5A-B**). Examination of tumors at the endpoint of tumor growth (15-20 days after IT *Lm* injection, late) also revealed a significant increase in the infiltration of CD8 T cells, but not a significant increase in CD4 T cells, in tumors of mice treated with IV+IT *Lm* compared to other groups (**Figures 5A-C**, **S5A-B**). Intratumoral regulatory T cells (Tregs) that are known to be immunosuppressive in the TME, were not significantly reduced with the IV+IT regimen compared to IV or IT alone, although they were reduced compared to the PBS treated mice (**Figures S5C-D**)^17,34–36,70–72^. Next, we assessed the functionality of the CD8 T cells by IFNγ and TNFα production 8 days after IT *Lm* or PBS injection. Here, the IV+IT *Lm* dosing regimen significantly increased CD8 T cells production of IFNγ, but especially the dual production of IFNγ and TNFα compared to PBS, IT *Lm* only, or IV *Lm* only (**Figure 5D**).

**Figure 5.**
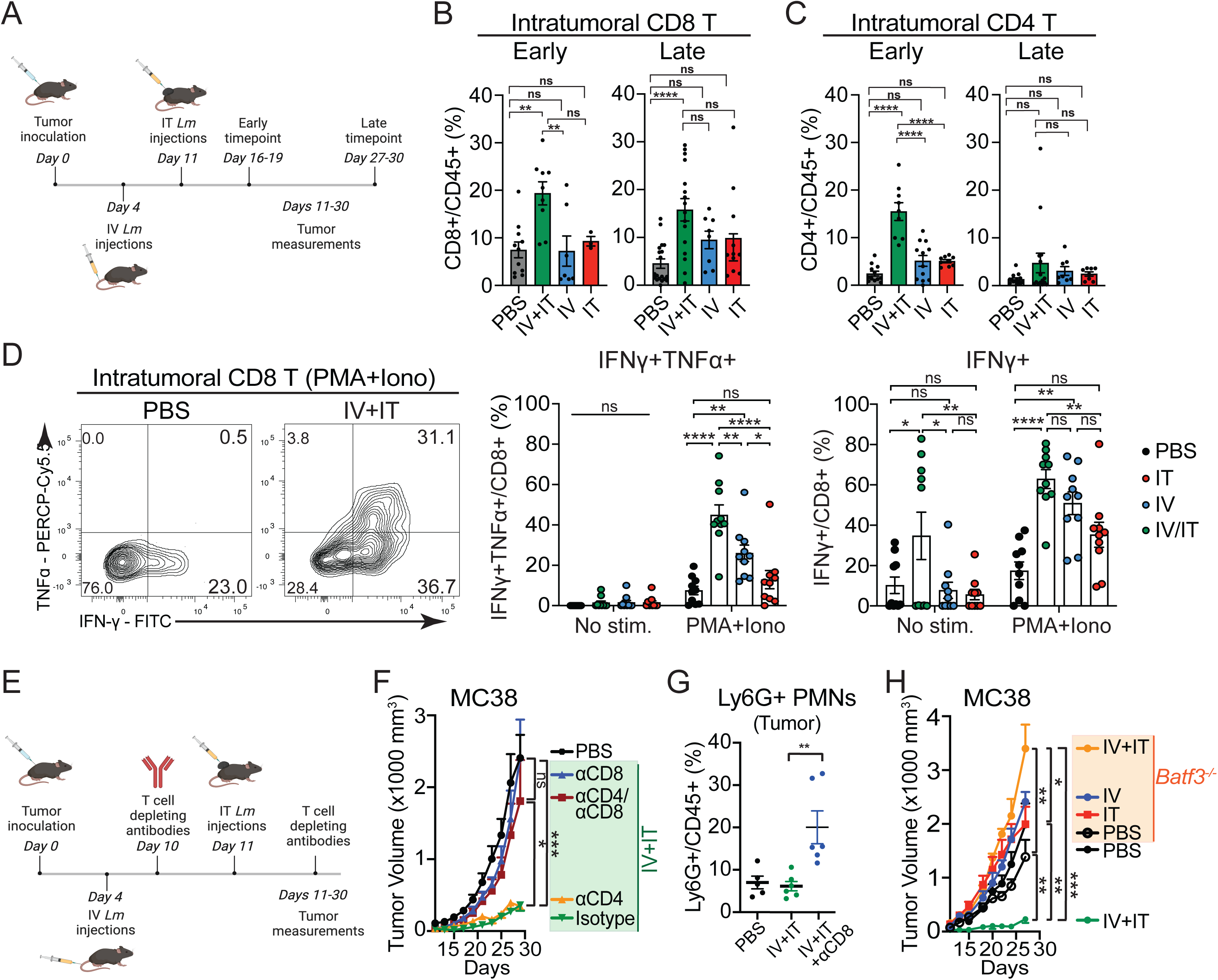
| CD8^+^ T cells are required for tumor control with IV+IT *Lm*. (A) Experimental strategy for IV+IT *Lm* dosing regimen followed by early or late flow cytometry analysis. (B-C) Frequency of intratumoral CD8^+^ (B) or CD4^+^ (C) T cells of CD45^+^ cells from MC38 tumors at early (left) or late (right) timepoints as specified in (A) (results pooled from four experiments with 5 mice/group). (D) Representative flow plots for IFNγ and TNFα intracellular cytokine staining from intratumoral CD8 T cells 8 days after IT *Lm* or IT PBS administration in each *Lm* dosing regimen (results pooled from two experiments with 5 mice/group). (E) Experimental strategy for IV+IT *Lm* dosing regimen plus anti-CD8, anti-CD4, or both depleting antibodies beginning 10 days after MC38 tumor inoculation. (F) MC38 tumor growth with T cell depletion. (G) Frequency of intratumoral Ly6G^+^ PMNs of CD45^+^ cells in IV+IT *Lm* treated mice +/-anti-CD8 depleting antibodies at end of experiment from (F). (H) MC38 tumor growth in WT versus *Batf3^-/-^* mice treated with IV+IT *Lm*. For all plots, *P<0.05, **P<0.01, ***P<0.001 by student t-test (B-C, G), one-way ANOVA (D), or two-way ANOVA (F, H), mean ± s.e.m.

To test whether CD4^+^, CD8^+^, or both populations of T cells were required for tumor control, we used antibody-mediated depletion of the T cells beginning one day prior to IT *Lm* injection **(Figure 5E)**. Depletion of CD8 T cells abrogated the efficacy of the IV+IT *Lm* regimen in MC38 and B16F10 models, indicating that CD8 T cells were required for tumor control with IV+IT *Lm* **(Figure 5F and Figure S5E-5F)**. In addition, CD8 T cell depletion increased the frequency of Ly6G^+^ cells maintained in tumors of mice treated with IV+IT *Lm*, suggesting that the reduction in PMN-MDSCs with IV+IT *Lm* is likely the result of CD8 T cell killing of *Lm*-infected PMNs (**Figure 5G**). We also tested the IV+IT dosing regimen in *Batf3^-/-^* mice that lack type I dendritic cells (DC1s), which are the primary DCs responsible for cross-presenting *Listeria* antigens to CD8 T cells^73,74^. The IV+IT dosing regimen did not lead to tumor control in *Batf3^-/-^* mice **(Figure 5H and Figure S5G)**. Taken together, the above results suggested that IV+IT *Lm* therapy requires the activity of T cells, in particular CD8 T cells. However, whether CD8 T cells specific to *Listeria*, the tumor, or both were required for tumor control could not be distinguished.

### *Listeria*-specific CD8 T cells drive tumor control

To assess whether the CD8 T cells that mediate tumor control with IV+IT *Lm* treatment are specific to *Lm* antigens, we used *Lm* engineered to express ovalbumin (*Lm-OVA*) and MHC-I (H2-K^b^) tetramers loaded with SIINFEKL to track the anti-*Lm* response (SIIN/H2K^b^) (**Figures 6A** and **S6A**). We observed an increased frequency of SIIN/H2K^b^ CD8 T cells in tumors with IV alone and IV+IT *Lm*, but only a significant increase in the absolute number of these *Lm*-specific cells in tumors with IV+IT *Lm* (**Figure 6A**). To test whether anti-*Lm* CD8 T cells alone could replace IV *Lm* administration to mediate tumor control, we used CD8 T cells from OT-I mice, which have a TCR that recognizes the OVA peptide SIINFEKL presented on MHC I (SIIN/H2K^b^). Adoptive transfer of activated OT-I CD8 T cells into mice prior to tumor inoculation allowed for a pool of *Lm*(SIIN)-specific memory CD8 T cells to develop in recipient mice, analogous to the expansion of *Lm*-specific memory CD8 T cells in prophylactically IV *Lm*-treated mice (**Figure 4G-H**). When tumors became palpable at day 11, tumors were injected with PBS, *Lm*, or *Lm-OVA* (**Figure 6B**). Strikingly, only mice whose tumors were injected with *Lm-OVA* had delayed tumor growth, similar to mice that received a standard IV+IT *Lm* regimen (**Figurse 6B** and **S6B**). Even in the aggressive orthotopic KP sarcoma model that does not express tumor-specific antigens^75^, the transfer of *Lm*-specific CD8 T cells prior to tumor implantation and IT *Lm-OVA* treatment was sufficient to provide significant tumor control (**Figures 6C** and **S6C**). These results indicate that a CD8 T cell response specific to *Lm* antigens is sufficient to phenocopy the effect of IV *Lm* in the IV+IT *Lm* dosing regimen.

**Figure 6.**
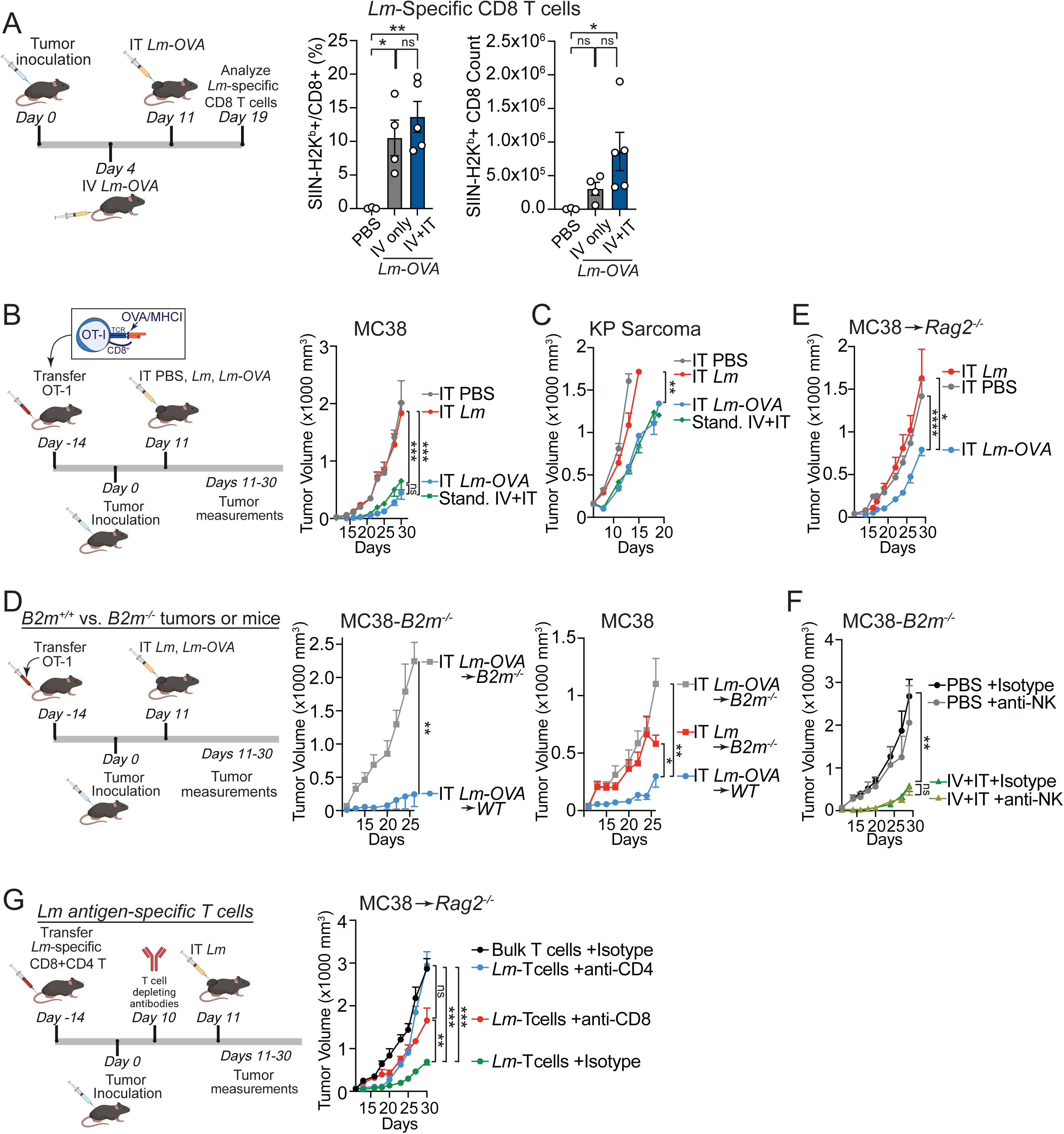
| Anti-*Listeria* CD8 immunity mediates tumor control. (A) Experimental design to analyze *Lm-*specific CD8 T cells after IV+IT *Lm* (right) frequency (middle) and total number (left) of *Lm*-specific (SIIN-H2K^b^ tetramer) CD8 T cells from MC38 tumors 8 days after IT *Lm-OVA* in IV+IT *Lm-OVA* regimen. OVA was expressed from *Lm* as a surrogate *Lm* antigen. (B) Experimental strategy to adoptively transfer *in vitro*-activated OT-I CD8 T cells in place of prophylactic IV *Lm* and MC38 tumors measured after IT PBS, *Lm*, or *Lm-OVA* (n= 5-6 mice/group from one of two experiments). (C) As in (B) but using an orthotopic KP sarcoma model (n= 5-6 mice/group from one of two independent experiments). (D) As in (B) but IT delivery compared MC38 tumors grown in WT or β2m^-/-^ mice (left) or MC38-*B2m^-/-^* tumors grown in WT or β2m^-/-^ mice (right) (n= 5-6 mice per group). (E) As in (B) but in *Rag2^-/-^* mice (n= 5-6 mice/group). (F) MC38-B2m^-/-^ tumor growth after IV+IT *Lm* or IV+IT PBS plus anti-NK1.1 beginning 10 days after MC38-B2m^-/-^ tumor inoculation (n= 5-6 mice/group). (G) Experimental strategy to adoptively transfer *in vitro*-activated *Lm*-specific CD8 and CD4 T cells in place of prophylactic IV *Lm* and MC38 tumors measured after IT PBS or *Lm* plus anti-CD8 or anti-CD4 beginning 10 days after MC38 tumor inoculation (n= 5-6 mice/group from one of two experiments). For all plots, *P<0.05, **P<0.01, ***P<0.001 by student t-test (A) or two-way ANOVA (B-G), mean ± s.e.m.

### *Listeria*-specific CD8 T cells control tumors without direct recognition of tumor cells

It has been reported that an alternatively attenuated strain of *Listeria* can directly infect cancer cells, and this could lead to the direct killing of tumor cells by *Listeria*-specific CD8 T cells^76^. While we did not find evidence of *Lm* infecting tumor cells using the *Lm-RFP* reporter strain **(Figures 2A-2D**), we wanted to test whether direct tumor cell killing by *Lm*-specific CD8 T cells was required for tumor control. To do this, we again transferred OT-I CD8 T cells prior to tumor development, but now tested their efficacy against *B2m^-/-^* (MHC I-deficient) MC38 tumors in either wildtype or *B2m^-/-^* mice, which lack MHC-I on all host cells (**Figure 6D and Figure S6D**). In this scenario, we could determine whether tumor control with IV+IT *Lm* required the recognition of infected tumor cells or host cells directly by OVA-specific CD8 T cells. In response to MC38-*B2m^-/-^* tumors, OT-I T cells mediated tumor control with IT *Lm-OVA*, but this only occurred in the setting of wildtype mice and not *B2m^-/-^* mice (**Figure 6D**). In the setting of wildtype MC38 tumors, MHC-I expression was required on host cells to mediate tumor control, as IT *Lm-OVA* was completely ineffective in *B2m^-/-^*mice despite the presence of OT-I T cells and MHC I expression on the tumor cells (**Figures 6D and S6E**). Furthermore, CFU analysis for live *Lm* also demonstrated that MHC I expression on host cells, but not tumor cells, was absolutely required for *Lm* clearance from the TME, in line with *Lm* persisting in host immune cells rather than tumor cells (**Figure S6F**). Thus, *Lm*-specific CD8 T cells require MHC I expression on host cells, but not tumor cells, to control tumors and bacterial load with the IV+IT *Lm* regimen.

To test whether the anti-*Lm* CD8 T cells alone were able to mediate tumor control, we injected *in vitro* activated OT-I cells into *Rag2*-deficient mice, which lack T and B cells and therefore cannot mount a direct antitumor CD8 T cell response. We then implanted MC38 tumors and IT injected PBS, *Lm* or *Lm-OVA* at day 11 when tumors became palpable. Here, tumor growth was still significantly reduced in *Rag-2^-/-^* mice that were IT injected with *Lm-OVA*, but not when injected with *Lm* that did not express OVA (**Figures 6E and Figure S6G**). Thus, tumor control could be mediated directly by anti-*Lm* CD8 T cells in the TME in the absence of additional host T cells.

To rule out a role for OT-I transgenic T cells and the recognition of OVA antigen in mediating tumor control, we also performed the standard IV+IT *Lm* regimen against MC38-*B2m^-/-^* tumors and still observed tumor control that did not require MHC I expression on the tumor cells (**Figure 6F and Figure S6H**). Furthermore, the lack of MHC I on tumor cells did not incite NK cell-mediated tumor control in response to missing self because IV+IT *Lm* controlled tumors equivalently with or without anti-NK1.1 depletion (**Figure 6F and Figure S6H**). Finally, we tested whether the transfer of T cells specific to natural *Lm* antigens could also drive tumor control by immunizing mice with *Lm*, expanding T cells from these mice *in vitro* with heat-killed *Lm*, and then prophylactically transferring bulk *Lm*-specific CD4 and CD8 T cells instead of OT-I T cells into *Rag2*-deficient mice. Here again, natural *Lm* antigen-specific T cells were able to control tumors in the absence of host T cells (**Figure 6G and Figure S6I**). Overall, these results indicate that T cells specific for *Lm* antigens that recognize non-tumor cells in the TME can mediate significant tumor control.

### *Listeria*-specific CD8 T cell killing and cytokine production drive tumor control

Next, we sought to determine how *Lm*-specific CD8 T cells, which do not target the cancer cells directly, mediate tumor control. First, we tested whether killing of non-tumor cells infected with *Lm* was required for the efficacy of the IV+IT *Lm* regimen by prophylactically transferring activated SIIN-specific CD8 T cells generated from WT versus *Prf1*-deficient (*Prf1^-/-^*) mice vaccinated with *Lm-OVA*. Mice were then inoculated with MC38 tumors, and at day 11 (when tumors were palpable), tumors were IT injected with *Lm* or *Lm-OVA* (**Figure 7A**). Tumor control was completely abolished when the transferred *Lm*-specific T cells lacked the pore-forming effector protein Perforin (**Figure 7B and Figure S7B**). Next, *Lm*-specific T cells that lacked the capacity to secrete IFN-γ (by harvesting SIIN-specific CD8 T cells generated from WT versus *Ifng^-/-^* mice vaccinated with *Lm-OVA*) also did not mediate tumor control (**Figure 7B and and Figure S7A**). Notably, the failure of *Prf1^-/-^* and *Ifng^-/-^* SIIN-specific CD8 T cells to mediate tumor control was not due to a failure of these cells to infiltrate the tumors, as an equivalent frequency of SIIN-specific CD8 T cells were found in tumors that received *WT*, *Prf1^-/-^*, or *Ifng^-/-^* SIIN-specific CD8 T cells (**Figure S7C)**. Since IV+IT *Lm* increased the production of IFN-γ and TNF-α from intratumoral CD8 T cells, we also tested whether secretion of TNF-α was required for the IV+IT *Lm* efficacy by using anti-TNF-α blocking antibodies. TNF-α blockade prevented tumor control with IV+IT *Lm* (**Figure 7C and Figure S7D**). Taken together, these results show that both the killing activity and cytokine production of *Lm*-specific CD8 T cells were required to mediate tumor control in the IV+IT *Lm* treatment regimen.

**Figure 7.**
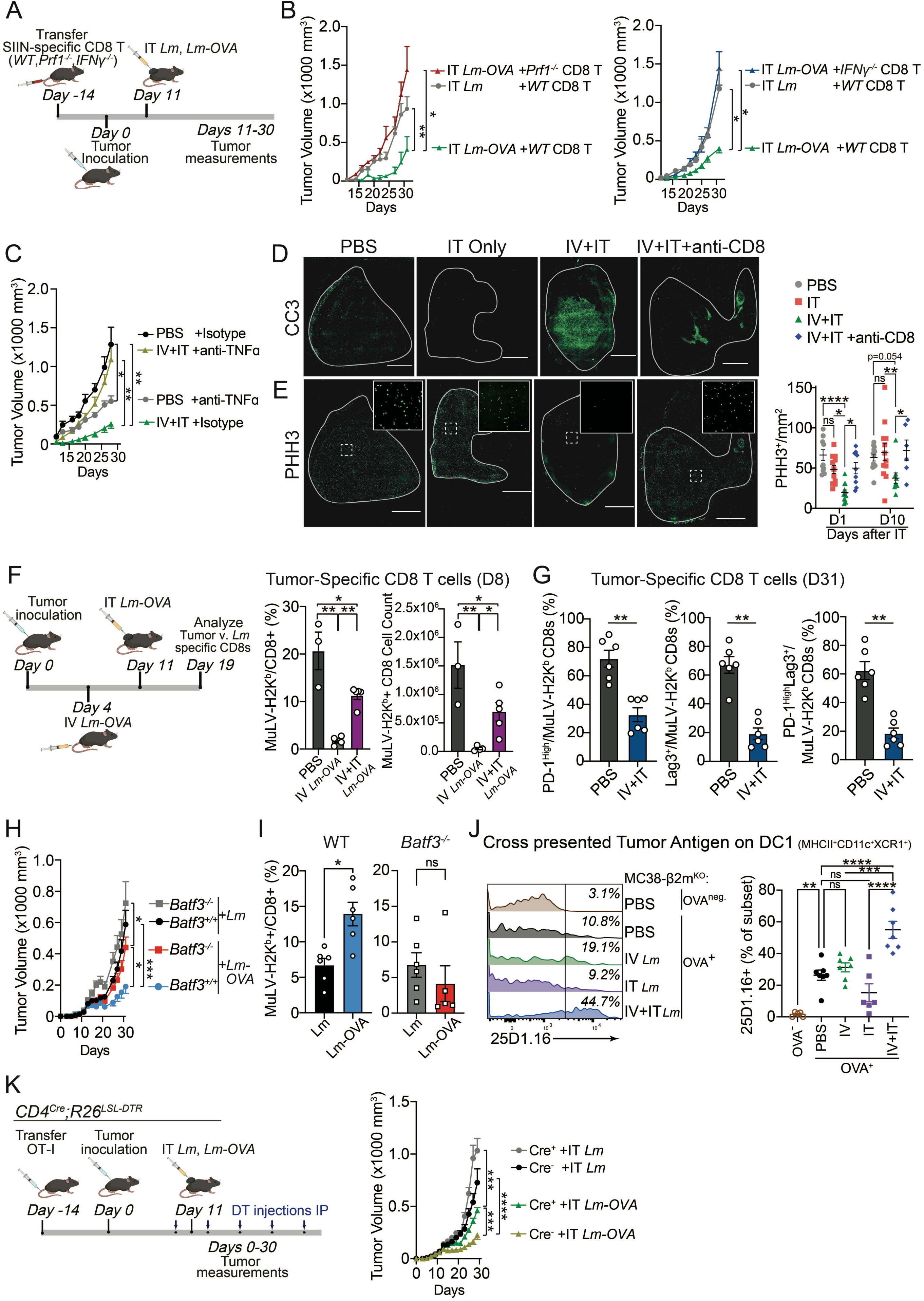
| Anti-*Listeria* CD8 immunity enhances antitumor responses. (A) Experimental strategy to adoptively transfer *in vitro* expanded SIIN-specific CD8 T cells from WT, *Prf1^-/-^, IFNγ^-/-^* mice immunized with *Lm-OVA* (see methods). (B) MC38 tumor growth after adoptive transfer of SIIN-specific CD8 T cells from WT, IFNγ*^-/-^*, or *Prf1^-/-^* mice followed by IT injection of PBS, *Lm*, or *Lm-OVA* at day 11 of tumor growth (n= 6 mice/group from one of two experiments). (C) MC38 tumor growth plus anti-TNFα beginning 10 days after MC38-B2m^-/-^ tumor inoculation (n= 5-6 mice/group from one of two experiments) (D) Representative images of Cleaved-Caspase 3 (CC3) (green) in MC38 tumors 24 hours post injection with IT PBS, IT *Lm*, or IV+IT *Lm* +/- anti-CD8 i.p. Scale bar = 2000μm. See Fig S7 for additional stains and day 10 images. (E) Representative images of phospho-Histone 3 (PHH3) (green) in MC38 tumors 24 hours post injection with IT PBS, IT *Lm*, IV+IT *Lm* +/- anti-CD8 i.p. and quantification of PHH3 staining in MC38 tumors in each group 1 day and 10 days after IT injections. Scale bar = 2000μm. Inset = high resolution images of boxed section of tissue. See Fig S7 for additional stains and day 10 images. (F) Experimental design for the analysis of frequency (left) and total number (right) of tumor*-*specific CD8 T cells (MuLV-H2K^b^ tetramer) 8 days after IT *Lm-OVA* in IV+IT *Lm-OVA* regimen compared to IV+IT PBS or IV *Lm-OVA* only. (G) Frequency of PD-1^High^ (left), Lag3^+^(center), and PD-1^High^Lag3^+^ double-positive (right) tumor-specific (MuLV-H2K^b^ tetramer) CD8 T cells from MC38 tumors 31 days after PBS or IV+IT *Lm*. (H) In vitro activated OT-I CD8 T cells were adoptively transferred into WT or *Batf3^-/-^*mice, MC38 tumors inoculated, and then tumors were injected IT on day 11 with *Lm* or *Lm-*OVA and tumor growth monitored (n= 5-6 mice per group pooled from two experiments). (I) Frequency of tumor-specific (MuLV-H2K^b^ tetramer) CD8 T cells from tumors at end of experiment in (H). (J) Representative histogram plots and quantification of 25D1.16 staining (anti-SIIN-H2K^b^) on intratumoral cDC1s (MHCII^+^CD11c^+^XCR1^+^) in OVA-expressing *B2m*-deficient MC38 tumors 24 hours post IT PBS, IT *Lm*, or IV+IT *Lm*. (K) Adoptive transfer of *Lm*-specific OT-I T cells (as in Figure 6B) was done into T cell depleter mice (*CD4-Cre; R26^LSL-DTR^*) that were treated with DT beginning 10 days after MC38 tumor inoculation and tumor growth was monitored (n= 5-6 mice per group pooled from two experiments). For all plots, *P<0.05, **P<0.01, ***P<0.001 by student t-test (F-G, I) or two-way ANOVA (B-C, H, J), mean ± s.e.m.

### *Listeria*-specific CD8 T cells enhance antitumor CD8 T cell responses

To determine how the IV+IT *Lm* regimen controlled tumors, we performed immunofluorescence analysis on tumors one and ten days after IT *Lm* (with prior IV *Lm*) or IT PBS (with prior IV PBS). We used anti-cleaved caspase 3 (CC3) staining to monitor the induction of apoptosis, as the requirement for Perforin in anti-*Lm* CD8 T cells indicated an important killing activity for the effectiveness of IV+IT *Lm* (**Figure 7B**). Indeed, the IV+IT *Lm* regimen induced a large amount of CC3 staining throughout the tumors one day after IT *Lm* injection, and this increased CC3 staining was markedly reduced by depleting CD8 T cells prior to IT injection (**Figure 7D**). The increase in CC3 staining was transient, however, as no differences were noted ten days after IT *Lm* injection (**Figure S7E**). In addition to their potential role in promoting cell killing, the requirement for IFN-γ and TNF-α may also be to slow tumor cell proliferation directly^77^. We used anti-phospho-histone H3 (PHH3) staining to monitor tumor cells in mitosis as a marker of cell proliferation and observed a nearly complete loss of PHH3 staining in tumor cells after IV+IT *Lm*, indicative of a near complete inhibition of cell cycle progression (**Figure 7E**). Notably, this reduced proliferation was still observed ten days after IT injection, though less robustly (**Figure S7E**). Thus, increased apoptosis and reduced cell proliferation likely contribute to the elimination and reduced growth of IV+IT *Lm*-treated tumors.

The massive induction of cell death upon IV+IT *Lm* may increase the phenomenon of epitope spreading, wherein death of tumor cells leads to the increased release and presentation of tumor antigens^78,79^. Thus, IV+IT *Lm*, while effective in controlling tumor progression with *Lm*-specific CD8 T cells alone, may also enhance the host T cell response directly against tumors. Notably, tumor control with IV+IT *Lm* was better in WT compared to *Rag-2^-/-^* mice, suggesting that host tumor-specific T cells may contribute to the efficacy of IV+IT Lm. (**Figures 6B versus 6E**). To assess the tumor-specific CD8 T cell response with IV+IT *Lm*, we used MHC I tetramers loaded with a tumor-derived peptide from the endogenous Murine Leukemia retrovirus, p15E (**Figure S7F)**^71,72^. Interestingly, while there was a decrease in the frequency and number of tumor-reactive (MuLV/H2K^b^ tetramer^+^) CD8 T cells in mice receiving the IV+IT *Lm-OVA* compared to PBS treated mice 8 days after IT injections, the frequency and number of these tumor-specific CD8 T cells was significantly increased compared to mice that only received IV *Lm-OVA* (**Figure 7F**). Furthermore, at later time points after IT injection (20 days), the tumor-specific T cells exhibited a phenotype of reduced T cell exhaustion (reduced PD1^+^Lag-3^+^ double and single positive cells) (**Figures 7G and S7G**). Thus, while IV *Lm* reduces tumor-specific T cell frequency and numbers, tumor-specific T cells are recalled to tumors following IV+IT *Lm* and retain a less exhausted phenotype throughout tumor growth.

Next we tested the importance of cross-presentation by type I dendritic cells (DC1), as we hypothesized that if anti-*Lm* CD8 T cells promoted tumor-specific CD8 T cell control, this would be dependent on DC1 tumor antigen cross-presentation^80,81^. We already observed that IV+IT *Lm* tumor control was completely abrogated in *Batf3^-/-^* mice (**Figure 5H**), likely due to the inability of mice to mount an anti-*Lm* CD8 T cell response after IV *Lm*^74,82^. Therefore, we used the transfer of activated OT-I CD8 T cells into *Batf3^-/-^* mice and then only IT injected *Lm* or *Lm-OVA*. We reasoned that this would establish whether cross-presentation after IT *Lm* was required to enhance that antitumor CD8 T cell response and increase tumor control. First, tumor control was only observed with IT *Lm-OVA* and not *Lm* as observed previously. As might be expected, *Batf3^-/-^* mice had significantly reduced tumor control compared to *Batf3^+/+^* mice with IT *Lm-OVA* **(Figure 7H and S7H)**. However, *Batf3^-/-^*mice treated with IT *Lm-OVA* also had significantly better tumor control than *Batf3^-/-^* mice treated with IT *Lm*, indicating that the OT-I CD8 T cells targeting *Lm* provided significant tumor control in the absence of DC1s (**Figures 5H versus 7H and S7H**). Most importantly, the frequency of tumor-specific (MuLV/H2K^b^ tetramer^+^) CD8 T cells was significantly increased in IT *Lm-OVA-* compared to IT *Lm*-treated mice, and this increase was totally dependent on the presence of Baft3^+^ DC1s in mice (**Figure 7I**). Finally, using a method we have employed previously to look directly at tumor antigen cross presentation on DC1s using OVA-expressing tumor cells and the 25D1.16 monoclonal antibody that detects SIINFEKL peptide from OVA presented on MHC I (H2-K^b^)^17,34–36^, we observed increased cross-presented tumor antigen on DC1s only with IV+IT *Lm*, but not with IV or IT only *Lm* administrations (**Figure 7J**). These results support the hypothesis that the *Lm*-specific CD8 T cells are able to enhance the tumor-specific CD8 T cell response by promoting the cross presentation of tumor antigens on Batf3^+^ DC1s, which may be particularly important for sustained control of tumors at later time points.

To directly test whether antitumor CD8 T cells also play a role in tumor control in our IV+IT *Lm* dosing regimen, we monitored tumor growth in the presence or absence of host T cells in the context of adoptive OT-1 T cell transfer to surrogate *Lm-*specific CD8 T cells. To do this, we generated a T cell depleter mouse model by combining *CD4-Cre* and *R26^LSL-DTR^* alleles^82^. Because all host T cells upregulate CD4 during thymic development, both mature CD4 and CD8 T cells permanently express the diphtheria toxin receptor and are depleted with the addition of diphtheria toxin (DT) (**Figure S7I)**^83^. Into these mice, we adoptively transferred *in vitro* activated OT-I CD8 T cells, transplanted MC38 tumors two weeks later, and then began DT treatment one day prior to IT injection of *Lm* or *Lm-OVA*. In the setting of tumor control with IT *Lm-OVA*, we now observed that the depletion of host T cells by DT treatment in *CD4-Cre^+^*(host T cells depleted) compared to *CD4-Cre^-^* (host T cells not depleted) mice significantly reduced tumor control (**Figures 7K and S7J)**. However, as expected, mice were still able to significantly control tumors without host T cells when treated with *IT Lm-OVA* compared to IT *Lm* (comparing *CD4-Cre^+^* mouse groups), indicating that *Lm*-specific OT-I CD8 T cells alone do provide significant tumor control (**Figures 7K and S7K**). Altogether, these data demonstrate that, in addition to *Lm*-specific CD8 T cells controlling tumors directly, they also enhance the tumor-specific CD8 T cell response by promoting the expansion and function of these CD8 T cells to increase tumor control.

## Discussion

In this study, we report two major findings. First, we showed that direct intratumoral injection of an attenuated strain of *Lm* caused increased tumor growth. This was due to *Lm*-mediated recruitment of PMNs that are converted within tumors into immunosuppressive MDSCs, which act as permissive cells for bacterial survival while also suppressing antitumor CD8 T cells. Second, we showed that IT *Lm* tumor growth promotion could be converted to tumor control by prior IV *Lm* immunization, which generated anti-*Listeria* CD8 T cells whose activity alone could mediate cancer protection. Thus, depending on the context of intratumoral seeding of *Lm* into a tumor, *Listeria* can either promote or inhibit cancer progression. These results reveal new insights into the biology of *Listeria* colonization of tumors in immune competent mice as well as the potential for anti-*Lm*-specific CD8 T cells to contribute to cancer control. These findings redefine the therapeutic mechanisms of action of bacterial cancer therapy and reveal the potential of immunological targeting of bacteria that colonize tumors to treat cancer.

The *ΔactA* attenuated *Lm* strain used in this study selectively colonized and persisted in tumors when injected IT or IV. Other studies have also reported short-term persistence of bacteria in tumors, either growing intracellularly (e.g. *Salmonella*) and/or extracellularly (e.g. *Clostridium*, *Vibrio. cholerae*, *E. coli)*, which in some cases had direct tumoricidal effects^5,84,85^. *Clostridium novyi* administered intravenously led to sporulation in the avascular regions of tumors that directly caused cancer cell death^13,86^. Longer term protection from cancer with *Clostridium* occurred in ∼30% of mice and depended on the generation of CD8 T cells, though the specificity of the CD8 T cells, against bacteria or tumor, was never directly tested^13,86^*. E. coli* engineered to express tumor antigens, the pore forming protein α-hemolysin, or L-arginine to promote CD8 T cell function, both led to improved tumor control upon bacterial colonization of tumors^21,87,88^. The prevalence of bacteria in tumors has been attributed to the presence of nutrients or an environment in tumors that is conducive to bacterial survival and growth^12,13,89–91^. *Salmonella typhimurium* engineered to infiltrate tumors by selective auxotrophy was effective at controlling several types of mouse tumors^12,89,92^. The lack of documentation of *Lm* persistence in tumors could be the result of using alternative attenuated strains of *Listeria* or failing to look for *Lm* persistence in previous studies. However, in the case of *ΔactA*-attenuated *Listeria* colonization in this study, we hypothesize that bacterial migration via the bloodstream is stochastic, but upon arrival in tumors, *Listeria* is protected from immune clearance due to the immunosuppressive environment created in tumors that generates PMN-MDSCs that permit the intracellular growth of *Lm*^5,93^.

PMN permissiveness for cytosolic *Lm* was strongly correlated with a CD14^+^ and PD-L1^+^ phenotype of the PMNs, markers associated with a MDSC phenotype. We also found that this phenotype was promoted by tumor derived factors, since culture of PMNs with tumor cell supernatants was sufficient to induce their expression of CD14 and PD-L1. In our system, however, depletion of these PMN-MDSCs did not prevent *Lm* persistence in tumors, but did prevent the accelerated tumor growth observed with IT *Lm* alone. This suggests that the TME prevents *Lm* clearance in multiple phagocytic immune cell types, but that recruitment and conversion of PMNs to MDSCs specifically promotes cancer progression. PMN-MDSCs and other myeloid cells have been found to promote tumor growth by several mechanisms, such as inhibiting T cell responses as well as by promoting vasculature formation to support tumor cell proliferation in glioblastoma, pancreatic, and several other mouse tumor models^35,38,64^. Using *in vitro* co-culture of FACS-purified intratumoral PMN-MDSCs with tumor cells and antitumor CD8 T cells, we showed that PMN-MDSCs could inhibit cytokine production from CD8 T cells responding to antigen recognition on tumor cells. Altogether, we uncovered an unexpected consequence of *Lm* seeding and colonization of tumors, wherein bacterial colonization of tumors by *Lm* does not lead to direct tumoricidal effects, but rather, recruits immune cells that are subsequently converted into immunosuppressive cells that promote tumor growth. However, microbes have also been shown to directly promote immune tolerance, and so it is possible that *Lm* directly promotes tolerance in the TME and its clearance with the IV+IT regimen allows for stronger tumor control^94^.

Attenuated *Listeria* vaccines generate memory CD8 and CD4 T cells that provide robust and long-lasting adaptive immunity^24,65^. We tested whether prior IV *Lm* administration to generate anti-*Lm* CD8 T cells would impact the immunosuppressive TME and promote tumor control with a secondary IT *Lm* dose. In this setting, IT *Lm* led to tumor control across multiple tumor models, including an aggressive orthotopic KP model of sarcoma. Tumor control was also observed if mice were prophylactically vaccinated prior to tumor inoculation, ruling out a role for tumor colonization with IV *Lm* in the efficacy of this IV+IT *Lm* regimen. Our results parallel previous approaches taking advantage of TDAP vaccination and using *Listeria* as a delivery system to introduce tetanus toxoid into tumors to elicit a CD4^+^ memory T cell response to eliminate tumor cells^76^. However, unlike that study, we found a clear requirement for CD8 T cells in mediating tumor control. Furthermore, in prior reports, it was shown that the *Listeria* infected the cancer cells directly to drive T cell recognition and control of cancer^76^. However, in our study we used a RFP reporter to identify cytosolic *Lm* and did not find evidence of direct infection of tumor cells. Nor was direct recognition of tumor cells by anti-*Lm* CD8 T cells required for tumor control, as MHC-I-deficient (*B2m*^-/-^) tumors were also controlled by the OT-I+IT *Lm-OVA* or the IV+IT *Lm* regimen. Furthermore, control of MHC-I-deficient tumors was not mediated by NK cells. However, MHC-I expression on non-tumor cells was essential for tumor control, as *B2m*-deficient mice receiving anti-*Lm* OT-I T cells were unable to control tumors after IT *Lm-OVA* administration.

Therefore, CD8 T cell recognition of host cells, presumably CD45^+^ immune cells that have taken up *Lm* in their cytosol, including PMN-MDSCs, was required for tumor control. The differences in mechanisms across studies could be the result of differences in the attenuated strains of *Listeria* used. Our study used an *ΔactA* mutation to attenuate the bacteria and prevent cell to cell spread, whereas others have used alternatively attenuated *Listeria* that retains a functional *actA* gene, thus allowing for the spread of *Listeria* to non-phagocytic cells, including tumor cells^17–20,23,76,91,95^. Overall, while direct tumor cell infection may support immune responses to tumors, our results indicate that tumor control by CD8 T cells does not necessitate bacterial spread into tumor cells.

To test the specificity of the CD8 T cells required for tumor control with IT *Lm*, we first tested the importance of anti-*Lm*-specific CD8 T cells by adoptive transfer of TCR transgenic (OT-I) CD8 T cells or polyclonal natural *Lm* antigen-specific T cells in place of IV *Lm* administration. We then used *Lm* IT delivery to show that T cells targeting *Lm* antigens, which were incapable of recognizing tumor antigens, could mediate tumor control. Tumor control with *Lm*-specific T cells even occurred after transfer into *Rag-2^-/-^* mice that could not generate any tumor-specific T cell responses, showing clearly that bacteria-directed T cells responding within the TME can control cancer. These results are contrary to the established dogma that an increase in tumor-specific CD8 T cells is required for the efficacy of bacterial-based cancer immunotherapies^96^. Of note, in clinical trials using BCG treatment for bladder cancer, patients with a positive skin tuberculosis test (PPD) due to previous vaccination with BCG, responded better to the BCG treatment and had longer recurrence-free survival than patients who did not establish pre-existing BCG-specific T cell immunity^97^.

To identify the functionalities of anti-*Lm* CD8 T cells required for tumor control, we transferred *Lm*-specific *Prf1^-/-^* or *Ifng^-/-^* CD8 T cells or used TNF-α blockade to show that killing activity, as well as IFN-γ and TNF-α production, were required for tumor control. We hypothesize that killing activity by CD8 T cells is required to eliminate PMN-MDSCs and the *Listeria* residing within them. This was evidenced by a high level of CC3-staining one day after IT *Lm* delivery exclusively in the IV+IT *Lm*-treated tumors that was dependent on CD8 T cells. IFN-γ and TNF-α production may be necessary to induce MHC I expression on these cells to mediate robust killing of *Lm*-infected cells, and/or it may act directly on tumor cells to reduce tumor cell proliferation^98–100^. In line with this, we observed a marked reduction in PHH3^+^ tumor cells with IV+IT *Lm* immediately following IT *Lm* delivery at one day and a sustained reduction in tumor cell proliferation for up to 10 days afterwards that was also dependent on CD8 T cells.

Finally, and perhaps most interestingly, IV+IT *Lm* enhanced the accessibility to tumor-derived antigens for cross presentation by cDC1s to tumor-specific CD8 T cells by an antigen spreading phenomenon^101–103^. This was evidenced by both increased tumor antigen presentation on DC1s and increased tumor-specific CD8 T cells infiltrating tumors with IV+IT *Lm* compared to IV *Lm* alone. In addition, tumor-specific CD8 T cells were less exhausted. We confirmed the importance of DC1s and tumor-specific T cells by genetic deficiency for DC1s (*Batf-3^-/-^*) or tumor-specific T cells (*CD4-Cre;R26^LSL-DTR^*, T depleter) and showing significant loss of tumor control in either scenario. However, the activity of anti-*Listeria* CD8 T cells were still paramount to tumor control, as their depletion completely restored rapid tumor growth. Thus, the anti-*Listeria* T cell response not only controls tumors indirectly, by promoting strong, localized cell death in the tumor and inhibiting tumor proliferation, but these activities also increase epitope spreading of tumor antigens, enhancing the tumor-specific CD8 T cell response to provide long-lasting systemic tumor immunity.

In conclusion, we have shown bacterial immunotherapy can have unintended negative outcomes by driving the accumulation of immunosuppressive cell types within tumors. However, the tumor promoting impact of *Lm* can be overcome by using a *Lm* dosing regimen that first generates anti-*Lm* CD8 T cells before direct intratumoral administration. The CD8 T cell response against *Lm* can not only remove the immunosuppressive cells from tumors, but also through direct cell killing of non-tumor cells in the TME, promote tumor control. Because the T cell responses against certain pathogens (i.e. *Listeria*, BCG, influenza, etc.) have been heavily studied and characterized, these findings have far-reaching consequences and may reveal new applications for existing therapeutics and vaccines. In addition, we show that methods to expand pathogen-specific T cells *ex vivo* for adoptive transfer of bacterial or viral antigen-specific T cells are effective and may be safer than attempting to generate memory T cells by active vaccination in potentially immunocompromised individuals, like cancer patients^104^. Collectively, our study demonstrates new insights into the inefficiencies of current bacterial immunotherapy approaches and reveals a tumor antigen-free, bacterial-based dosing regimen that can control cancer independently of its immunogenicity or the identification of tumor neoantigens. Most importantly, the recent findings that many tumors are colonized by various bacterial species opens the door to a new alternative strategy to modify the tumor microenvironment and treat cancer, by targeting intratumoral bacteria.

### Limitations of the study

This study only used one attenuated strain of *Listeria monocytogenes* for all investigations and while we hypothesize that the findings will be generalizable to other bacterial species, or even other attenuated strains of *Listeria*, this remains to be tested. Future studies will also need to test what specific bacterial attributes of *Listeria* are required to promote tumor control - such as the capacity for intracellular invasion or the capacity to activate specific innate pattern recognition receptors, which will shed light on the broad applicability of these findings to other bacterial species. Next, while we showed that *Lm* colonization of a non-subcutaneous tumor model (PDAC) also occurred, further investigations to test whether the IV+IT (or IV+IV) *Lm* therapeutic strategy is also effective against internally arising epithelial cancers need to be performed. Finally, the findings here rely solely on mouse cancer models. Future studies should examine whether human tumors will yield similar results, such as with humanized mouse models or the examination of human tumor samples, especially from patients who have been treated with *Listeria* vaccines.

## Acknowledgments

We thank Russell Vance, Gregory Barton, David Raulet, and Thomas Dubensky for reviewing the manuscript and providing feedback; Hector Nolla, Alma Valleros, Kartoosh Heydari and Anita Wong Lin of the UC Berkeley Cancer Research Laboratory Flow Cytometry Facility. This research was supported by 1DP2CA247830-01 from the National Institutes of Health (NCI) and C23CR5612 from the UC CRCC Cancer Research Coordinating Committee. M.D. is a Pew-Stewart Scholar and a St. Baldrick’s Scholar with generous support from Hope with Hazel. J.G.C is a HHMI Gilliam Fellow.

## Author Contributions

Conceptualization and Methodology, M.D., D.A.P., J.G.C., S.F., T.F.C; Investigation, J.G.C., S.F., T.F.C., J.W., D.G.V., J.Y., N.F.H., E.W., and M.D.; Writing – Original Draft, M.D., J.G.C., S.F.; Writing – Review & Editing, M.D., D.A.P., J.G.C., S.F., T.F.C; Supervision, M.D., D.A.P., J.G.C., S.F.; Funding Acquisition, M.D., D.A.P.

## Declaration of Interests

D.A.P. is on the board of directors for Laguna Therapeutics.

## STAR Methods

### Resource Availability

#### Lead Contact

Further information and requests for resources and reagents should be directed to and will be fulfilled by the lead contact, Michel DuPage.

#### Materials availability

*Listeria* monocytogenes strains used in the study will be made available from the lead contact upon request.

#### Data and code availability

Any additional information required to reanalyze the data reported in this paper is available from the lead contact upon request.

### Experimental models and subject details

#### Animal studies

C57BL/6J wildtype mice were obtained from Jackson laboratories (JAX:000664) and bred in house. OT-1 transgenic mice were obtained Taconic (Catalog#: 2334) and bred in house. Prf1^-/-^ mice were a gift from the Stanley lab at the University of California, Berkeley (JAX:000274). Rag2^-/-^ (JAX:008449) and β*2m2m^-/^*^-^ (JAX:002087) mice were a gift from the Raulet lab at the University of California, Berkeley. CD4^Cre^ mice were obtained from Jackson laboratories (JAX:022071) and bred in house. R26^DTR^ mice were obtained from Jackson laboratories (JAX:008040) and bred in house. Sarcoma cell lines were generated in *Kras^LSL-G12D/+^;p53^fl/fl^ mice* by intramuscular injection of the left hind limb with replication-incompetent lentiviruses expressing Cre recombinase as reported previously, harvested, and cultured^75^. For tumor studies, syngeneic C57BL/6J mice were inoculated with 5.0×10^5^ MC38 or 2.0×10^5^ B16F10 cells in PBS subcutaneously. KP sarcoma cells were inoculated into C57BL/6J mice with 5.0×10^5^ cells by intramuscular injection of the right hind limb. For diphtheria toxin (DT) treatment for T cell depletions, we injected 1μg every 2 days by i.p. Tumor measurements were performed blindly across the entire experiment by a single operator measuring three dimensions of the tumor with calipers three times per week. All the experiments were conducted according to the Institutional Animal Care and Use Committee guidelines of the University of California, Berkeley.

#### Cell lines

MC38 and B16F10 cell lines were kindly provided by Dr. Jeff Bluestone’s lab^105^. MC38-β2m^-/-^ cell line was kindly provided by Dr. David Raulet’s lab^47^. All cell lines were maintained in DMEM (GIBCO) supplemented with 10% FBS, sodium pyruvate (GIBCO), 10mM HEPES (GIBCO), and penicillin-streptomycin (GIBCO). Tumor cells were grown at 37℃ with 5% CO_2_.

### Method Details

#### Listeria monocytogenes strains

All strains of *L. monocytogenes* were derived from the wild-type 10403S strain. The *Lm* constructs were based on an attenuating deletion of the actA gene (*ΔactA*). *Lm-OVA* expresses an LLO-OVA fusion encompassing amino acids 1-441 of Listeriolysin O (LLO) and OVA cloned in frame of LLO1-441 under the control of the LLO promoter in the pPL2 integration vector^106^. *Lm-TagRFP* expresses a secreted TagRFP protein under the control of the *actA* promoter using the pPL2 integration vector^52^. All strains were cultured in filter-sterilized nutrient-rich Brain Heart Infusion (BHI) media (BD Biosciences) containing 200 μg/mL streptomycin (Sigma-Aldrich).

#### Intravenous and Intratumoral *Listeria* infection

Overnight cultures were grown in BHI + 200 μg/mL streptomycin at 30℃. The following day, bacteria were grown to logarithmic phase by diluting the overnight culture in fresh BHI + 200 μg/mL streptomycin and culturing at 37℃ shaking. Log-phase bacteria were washed and frozen in 9% glycerol/PBS. For intravenous infections, frozen stocks were diluted in PBS to infect via the tail vein with 1 x 10^6^ CFU log-phase bacteria. For intravenous infections, frozen stocks were diluted in PBS to infect via intratumoral infections at 5 x 10^7^ CFU log-phase bacteria. The mice were euthanized 8-20 days after intratumoral injections and organs were collected for flow analysis.

#### Colony forming unit assays from tissues

Tissues were collected in 0.1% NP40 buffer diluted in PBS. Organs were homogenized and serially diluted on non-TC treated 96-well plates (Genesee). Serial dilutions were plated on BHI + 200 μg/mL streptomycin plates. Plates were incubated overnight at 37℃.

#### CIN/Primary bone-marrow derived PMN Maintenance and Differentiation

Cas9+ ER-Hoxb8 conditionally immortalized neutrophils (CINs) were a gift from the Stanley lab at the University of California, Berkeley^61^. CINs were expanded as progenitors in Optimem with 10% FBS, 1% L-glutamine, 30μM 2-mercaptoethanol, and 1% Stem-cell factor (SCF)-producing CHO cell supernatant (CIN Media) that also contained 1μM beta-estradiol in non-TC treated flasks (Genesee) maintaining a concentration less than 1×10^6^ cells/mL. Upon reaching the desired quantity, non-adherent progenitors were harvested, washed twice in cold PBS 1x, and plated in non-TC treated flasks containing CIN media lacking beta-estradiol. To generate PMNs, CINs were differentiated until Day 6 in CIN Media + 5ng/mL GM-CSF. To generate PMN-MDSCs, CINs were differentiated in CIN media + 5ng/mL GM-CSF, 40ng/mL IL-6, and 40ng/mL GM-CSF. For primary bone marrow PMNs and PMN-MDSCs, cells were isolated and cultured as described^63^. To generate PMNs-MDSCs, cells were cultured in 40ng/mL IL-6 and 40ng/mL GM-CSF1 day post isolation for 6 days.

#### In vitro infections

Cells were infected with *Lm-TagRFP* at an MOI that resulted in 30% of the cells being infected. At 1hr post infection, 50μg/mL Gentamicin was added to kill all extracellular Listeria. For flow analysis, cells were collected at time points indicated, stained, and fixed with 4% PFA.

#### Cancer Supernatant Assays

Cancer cell supernatant was generated via the expansion of MC38, B16F10, and 3T3 cell lines in TC-treated flasks in Optimem + 10% FBS, followed by collection of supernatant fluid when cells reached near ∼100% confluency. After filtration, all supernatants were frozen down at -20°C. Surveying cancer supernatants involved a 24h incubation period where cells were either re-seeded into TC-treated 24 well plates or non-TC treated flasks in fresh CIN media with a 50:50 ratio of MC38, B16F10, or 3T3 supernatant. Cells were collected at 24h, stained, fixed with 4% PFA, and analyzed by flow cytometry.

#### Tissue Collection and preparation for Flow cytometry

Flow cytometry was performed on an BD LSR Fortessa X20 (BD Biosciences), CyTEK Aurora (CyTEK Biosciences) or LSRFortessa (BD Biosciences) and datasets were analyzed using FlowJo software (Tree Star). Single cell suspensions were prepared in ice-cold FACS buffer (PBS with 2mM EDTA and 1% BS) and subjected to red blood cell lysis using ACK buffer (150mM NH4Cl, 10mM KHCO3, 0.1mM Na2EDTA, pH7.3). Dead cells were stained with Live/Dead Fixable Blue or Aqua Dead Cell Stain kit (Molecular Probes) in PBS at 4℃. Cell surface antigens were stained at 4℃ using a mixture of fluorophore-conjugated antibodies. Surface marker stains for murine samples were carried out with anti-mouse CD3 (17A2, BioLegend), anti-mouse CD4 (RM4-5, BioLegend), anti-mouse CD8a (53-6.7, BioLegend), anti-mouse, CD44 (IM7, BioLeged), anti-mouse CD45 (30-F11, BioLegend), anti-H-2Kb MuLV p15E Tetramer-KSPWFTTL (MBL), anti-H-Kb-A2/SIINFEKEL tetramer (NIH tetramer core), anti-IAb/NEKYAQAYPNVS tetramer (NIH tetramer core) in PBS, 0.5% BSA. Cells were fixed using the eBioscience Foxp3/Transcription Factor staining buffer set (eBioscience), prior to intracellular staining. Intracellular staining was performed using anti-mouse Foxp3 (FJK-16S, eBioscience), anti-mouse TNF-α (MP6-XT22, BioLegend), anti-mouse IFN-γ (XMG1.2, eBioscience), at 4℃, according to manufacturer’s instructions. For Lm-TagRFP infected tissues, single cell suspensions were prepared as above, stained using a mixture of fluorophore-conjugated antibodies at 4℃, and fixed in 4% PFA. Cells analyzed by flow cytometry the following day to prevent signal loss from flourescent protein. Cells were resuspended in PBS and filtered through a 70-μm nylon mesh before data acquisition. Datasets were analyzed using FlowJo software (Tree Star).

#### Restimulation Assays

Resected tumors were minced to 1 mm^3^ fragments and digested in RPMI media supplemented with 4-(2-hydroxyethyl)-1-piperazi-neethanesulfonic acid (HEPES), 20 mg/mL DNase I (Roche), and 125 U/mL collagenase D (Roche) using an orbital shaker at 37℃. Cells from lymphoid organs were prepared by mechanical disruption pressing against a 70-μm nylon mesh. All the cell suspensions were passed through 40 μm filters before in vitro stimulation. Cytokine staining was performed with 3-5×10^6^ cells in Opti-MEM media supplemented with Brefeldin A (eBioscience) or 10 ng/mL phorbol 12-myristate 13-acetate (PMA) (Sigma), and 0.25 μM ionomycin (Sigma).

Fixation/permeabilization of cells was conducted for intracellular staining using the eBioscience Foxp3 fixation/permeabilization kit (BioLegend) or Tonbo Foxp3 / Transcription Factor Staining Buffer Kit.

#### Adoptive transfer experiments

For in vitro T cell culture, spleens and lymph nodes were collected from OT-1 transgenic mice or C57BL/6J wildtype (*WT*) or *Prf1^-/-^* previously vaccinated with 1.0×10^6^ CFU *Lm-OVA* 3-4 weeks prior. OT-1s or OVA-responsive CD8s were were activated with 1μg/mL SIINFEKL peptide in DMEM medium supplemented with 10% FBS, non-essential amino acids, sodium pyruvate, L-glutamine, HEPES, 55 μM β-ME and 200 IU/ml recombinant human IL-2 (TECIMTM, Hoffman-La Roche provided by NCI repository, Frederick National Laboratory for Cancer Research). T cells were transferred into WT or Rag2^-/-^ or *B2m^-/^*^-^ mice by intravenous injection two weeks prior to MC38 or KP sarcoma inoculations.

#### *In vivo* antibody-mediated cell depletion

For tumor progression studies, CD8 depletion was achieved by intraperitoneal injection of 200 μg per mouse of the anti-CD8 monoclonal antibody clone YST-169 (Leinco Technologies, Catalog # C2442) two days prior to tumor inoculation, followed by additional doses every 6 days thereafter. For CD4 depletion, intraperitoneal injection of 200 μg/mouse of clone GK1.5 (Leinco Technologies, Catalog: C1333) was done two days prior to tumor inoculation, followed by additional doses every 6 days thereafter.

PMN depletion was done by intraperitoneal injection of 200 μg per mouse of the anti-Ly6G monoclonal antibody clone IA8 (Leinco Technologies, Catalog: L280) twice prior to tumor inoculation, followed by additional doses every 2 days.

#### Immunofluorescence & Imaging Analysis

Tissue Sections were washed with PBS and blocked with tissue staining buffer plus 10% fetal bovine serum. Tissues were then stained with anti-Ly6G Alexa Fluor 488 antibody (Clone 1A8, Biolegend Catalog# 127624), anti-Phospho-Histone H3 (Ser10) antibody (Cell Signaling Catalog# 9701S, or anti-Cleaved Caspase-3 (Asp175) antibody (Clone 5A1E, Cell Signaling Catalog# 9664S) O/N at 4℃ in tissue staining buffer (PBS with 0.2% Triton X-100, 2% FBS) and washed with PBS. For sections stained with unconjugated primary antibodies, sections were incubated with Goat anti-Rabbit IgG (H+L) Cross-Adsorbed Secondary Antibody, Alexa Fluor™ 488 (Thermo Fisher Scientific, Catalog# A-11008) for 1 hour at RT followed by washes with PBS. Tissues were stained with Wheat Germ Agglutinin (WGA) Alexa Fluor 647 (Thermo Fisher Scientific, Catalog# W32466) for 45 mins at RT and washed with PBS. Stained tissue sections were mounted with Fluoromount-G Mounting Medium (Southern Biotech).

Images were taken using Zeiss LSM 780 NLO microscope and Zeiss Axioscan 7 or Axioscan Z1. Imaging analysis was performed using Zeiss Zen Lite Imaging Software or Fiji/ImageJ. Quantification of phH3 staining was performed by cropping full Axioscan images into 2 mm x 2 mm images from which 3 representative cropped images per tumor were chosen for counting. Individual PHH3+ cells in these sections were blocked out using the Fiji/ImageJ paintbrush tool until no PHH3+ cells remained visible. A click counter (Mousotron) was used to track the number of cells per section.

### Statistical Methods

p values were obtained from unpaired two-tailed Student’s tests for all statistical comparisons between comparisons between two groups, and data were displayed as mean ± SEMs. For multiple comparisons, one-way ANOVA was used. For tumor growth curves, two-way ANOVA was used with Sidak’s multiple comparisons test performed at each time point or by multiple regression analysis p values are denoted in figures by *p < 0.05, **p < 0.01, ***p < 0.001, and ****p < 0.0001.

**Figure S1 | Intravenous injection of *Lm* alone does not slow tumor growth.**

(A) Tumor growth curves for individual mice in 1B.

(B) Tumor growth curves for individual mice in 1C.

(C) Experimental design.

(D-E) C57BL/6J WT mice were injected with 5×10^5^ MC38 (B, data shown from n= 5 mice per group from one of three experiments) or 2×10^5^ B16F10 (C, data shown from n= 5 mice per group from one of two experiments) tumor cells and then 1×10^6^ *Lm* CFUs IV 4 days later and tumors measured.

(F) CFU analysis for *Lm* in MC38 tumors from mice injected IV with 1×10^6^ CFU 4 days after tumor inoculation showed seeding and persistence of *Lm* in tumors as in (B) (n=4-5 mice per time point).

For all plots, statistical analysis was done by two-way ANOVA, mean ± s.e.m.

**Figure S2 | Population analysis during intratumoral *Lm* infection in MC38 and B16F10 tumors.**

(A) Gating strategy for identifying immune cell subsets from single cell suspension of MC38 tumor.

(B) Proportion of immune cell types that were CD45^+^RFP^+^ from B16F10 tumors 24 hours after IT *Lm* (n= 4-5 mice/ group).

(C-D) Frequency of intratumoral immune cell populations as percent of all CD45^+^ cells 24 hours after IT PBS or *Lm* injection in MC38 (C) or B16F10 (D) tumors (n=4-5 mice/group).

(E) Frequency of Ly6G^+^ PMNs in spleen and B16F10 tumors 24 hours after IT PBS or IT *Lm* (n=4-5 mice/group).

(F) Frequency of intratumoral Ly6G^+^ PMNs that were RFP^+^ 24 hours after IT *Lm* in B16F10 tumors.

(G) Validation of anti-Ly6G staining on frozen tumor sections by testing sections from mice treated with an isotype control antibody or an anti-Gr1 antibody that depletes all Ly6G+ cells.

For all plots, mean ± s.e.m. and *P<0.05, **P<0.01, ***P<0.001 by student t-test (B, C, E, F, G, H).

**Figure S3 | PMN-MDSCs are permissive to *Lm* infection.**

(A) Frequency of CD14+ PMNs 24 hours after IT *Lm* in B16F10 tumors. Results from n= 4-5 mice/group

(B) Frequency of intratumoral CD14+Ly6G^+^ cells that were RFP^+^ 24 hours after IT PBS or IT *Lm*.

(C) Representative flow cytometry plots of the differentiation of primary bone marrow progenitors into CD14^+^ PMN-MDSCs *in vitro* by culturing with IL-6 and GM-CSF.

(D) Heterogeneous PMN-MDSCs differentiated from bone marrow progenitors as in (A) were infected with *Lm-RFP in vitro* and Tag-RFP^+^ cells were identified 4 or 8 hours later in all cells (Total PMNs), CD14^+^ cells, or CD14^-^ cells by flow cytometry.

(E) Tumor growth curves for individual mice in 3J.

(F) *Lm* CFUs recovered from tumors 20 days after IT *Lm* in mice treated with isotype control or anti-Ly6G antibodies as shown in Figure 3J.

(G) Proportion of immune cell types that were RFP^+^ from MC38 tumors treated with an isotype control (left) or anti-Ly6G antibody (right) 24 hours after IT *Lm*.

(H) Frequency of intratumoral immune cell populations as percent of all CD45^+^ cells 24 hours after IT *Lm* or IT PBS into MC38 tumors in mice pre-treated with isotype or anti-Ly6G (PMN-depleting) antibodies as shown in Figure 3I.

For all plots, *P<0.05, **P<0.01, ***P<0.001 by student t-test (A-B, F, H), or one-way ANOVA (D), mean ± s.d for *in vitro* experiments. Two-way ANOVA (J), mean ± s.e.m for *in vivo* experiments

**Figure S4 | IV+IT *Lm* is not impacted by PMN depletion and IV+IT regimen leads to *Lm* clearance.**

(A) Tumor growth curves for individual mice in 4B.

(B) Tumor growth curves for individual mice in 4C.

(C) Average (right) and individual tumor growth (left) for MC38 with IV+IV PBS versus IV+IV *Lm* versus IV+IT *Lm* regimen (data shown from n= 5-6 mice per group from one of two experiments).

(D) Average (right) and individual tumor growth (left) for MC38 tumor growth with PBS versus IV+IT *Lm* and with or without anti-Ly6G depleting antibody treatment beginning two days prior to IT *Lm* administration (n=4-6 mice per group).

(E) Tumor growth curves for individual mice in 4H.

(F) Tumor growth curves for individual mice in 4I.

For all plots, **P<0.05, **P<0.01, ***P<0.001 by two-way ANOVA (C-D), mean ± s.e.m.

**Figure S5 | IV+IT *Lm* dosing regimen reduces intratumoral Treg frequencies but does not impact T cell frequencies in the spleen.**

(A-B) Frequency of CD8^+^ (C) and CD4^+^ (D) T cells from spleens at early or late timepoints. Data pooled from four experiments with 5 mice/group.

(C) Frequency of intratumoral Foxp3^+^ Tregs of CD4^+^ cells from MC38 tumors at early (left) or late (right) timepoints as specified in Figure 5A.

(D) Frequency of Foxp3^+^ Tregs of CD4^+^ cells from spleens at early (left) or late (right) timepoints.

(E) Tumor growth curves for individual mice in 5F.

(F) Average (right) and individual tumor growth (left) for B16F10 with IV+IT *Lm* dosing regimen plus anti-CD8, anti-CD4, or both depleting antibodies beginning 10 days after B16F10 tumor inoculation.

(G) Tumor growth curves for individual mice in 5H.

For plots in S5A-S5D, timepoints are the same as what was described in Figure 5A. For all plots, *P<0.05, **P<0.01, ***P<0.001 by student t-test (A-D) or two-way ANOVA (F), mean ± s.e.m.

**Figure S6 | *Lm*-specific CD8 T cells are present in tumors independent of Prf1 genotype.**

(A) Representative flow plots for SIIN-H2K^b^ tetramer staining from tumors of mice treated with IV+IT PBS compared to IV+IT *Lm-OVA* 8 days after IT injections.

(B) Tumor growth curves for individual mice in 6B.

(C) Tumor growth curves for individual mice in 6C.

(D) Tumor growth curves for individual mice in 6D.

(E) Frequency of intratumoral OT-I CD8 T cells in mice 20 days after IT *Lm* administration from Figure 6D.

(F) CFU analysis for *Lm* in tumors from Figure 6D.

(G) Tumor growth curves for individual mice in 6E.

(H) Tumor growth curves for individual mice in 6F.

(I) Tumor growth curves for individual mice in 6G.

For all plots, *P<0.05, **P<0.01, ***P<0.001 by student t-test (E-F), mean ± s.e.m.

**Figure S7 | *Antitumor T cell immunity enhancement is dependent on Listeria-specific direct killing and cytokine secretion.***

(A-B) Tumor growth curves for individual mice in 7B.

(C) Frequency of intratumoral SIIN-specific CD8 T cells in mice 20 days after IT *Lm* administration from experiments depicted in Figures 6J-K (pooled from two independent experiments)

(D) Tumor growth curves for individual mice in 7C.

(E) Images of MC38 tumors 24hrs post injection with IT PBS, IT *Lm*, or given IV+IT dosing regimen +/- anti-CD8 i.p: phospho-Histone 3 (PHH3) or cleaved-caspase 3 (CC3) (green), WGA (red) and DAPI (blue). Scale bar = 2000μm.

(F) Images of MC38 tumors 10 days post injection with IT PBS, IT *Lm*, or given IV+IT dosing regimen +/- anti-CD8 i.p: phospho-Histone 3 (PHH3) or cleaved-caspase 3 (CC3) (green), WGA (red) and DAPI (blue). Scale bar = 2000μm.

(G) Representative flow plots for MuLV-H2K^b^ tetramer staining in lymph nodes from naive mice compared to MC38 tumor-bearing mice 19 days post tumor inoculation.

(H) Representative flow plots for PD-1+Lag3+ MuLV-H2K^b^ CD8 T cells for 7G.

(I) Tumor growth curves for individual mice in 7H.

(J) Depletion of T cells in T cell depleter mice (CD4^Cre^; R26^LSL-DTR^) in non-tumor bearing mice. Mice were injected twice with 1μg DT i.p. every two days and inguinal lymph nodes and spleen collected for flow analysis.

(K) Tumor growth curves for individual mice in 7K.

(L) Frequency of intratumoral tumor-specific (MuLV-H2K^b^ tetramer) CD8 T cells (right) and OT-I CD8 T (left) cells in mice 20 days after IT *Lm* administration from Figure 7J. For all plots, *P<0.05, **P<0.01, ***P<0.001 by student t-test (C,M) or two-way ANOVA (H), mean ± s.e.m.

## Notes

### Summary of Updates

Update to Figures 1,2,3, and 6 with the addition of figure 7 not found in the first version of the manuscript. In addition, supplemental figures have been updated as well.

