## Supplemental Figures S1-S7 for "Cellular mechanisms underlying beneficial versus detrimental effects of bacterial antitumor immunotherapy"

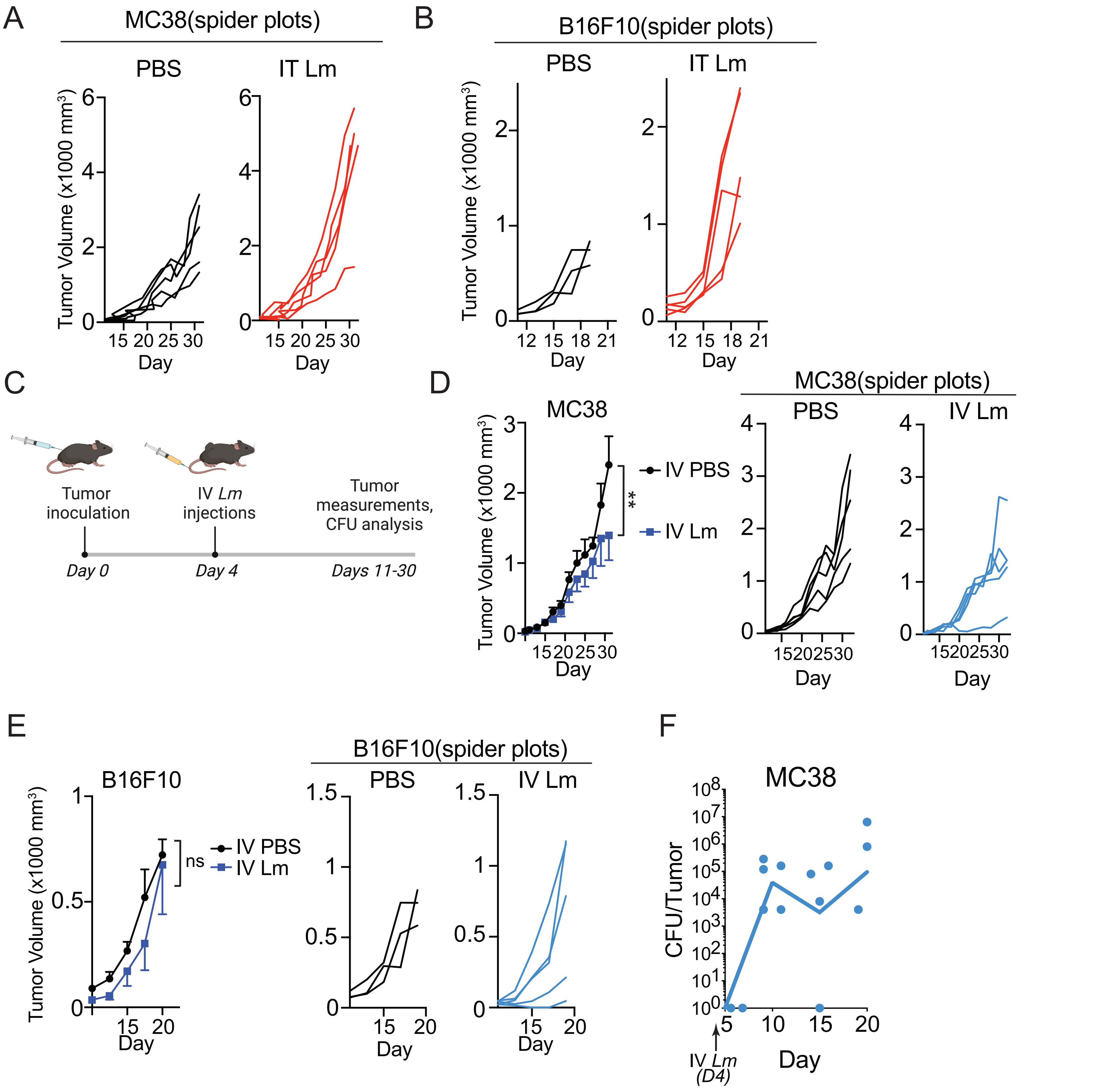


Garcia Castillo, J., Campbell, T., & Fernandez, S. et al. Figure S1.

**Figure S1 | Intravenous injection of *Lm* alone does not slow tumor growth.**
(A) Tumor growth curves for individual mice in 1B.
(B) Tumor growth curves for individual mice in 1C.
(C) Experimental design.
(D-E) C57BL/6J WT mice were injected with $5x10^5$ MC38 (B, data shown from n= 5 mice per group from one of three experiments) or $2x10^5$ B16F10 (C, data shown from n= 5 mice per group from one of two experiments) tumor cells and then $1x10^6$ *Lm* CFUs IV 4 days later and tumors measured.
(F) CFU analysis for *Lm* in MC38 tumors from mice injected IV with $1x10^6$ CFU 4 days after tumor inoculation showed seeding and persistence of *Lm* in tumors as in (B) (n=4-5 mice per time point).
For all plots, statistical analysis was done by two-way ANOVA, mean ± s.e.m.

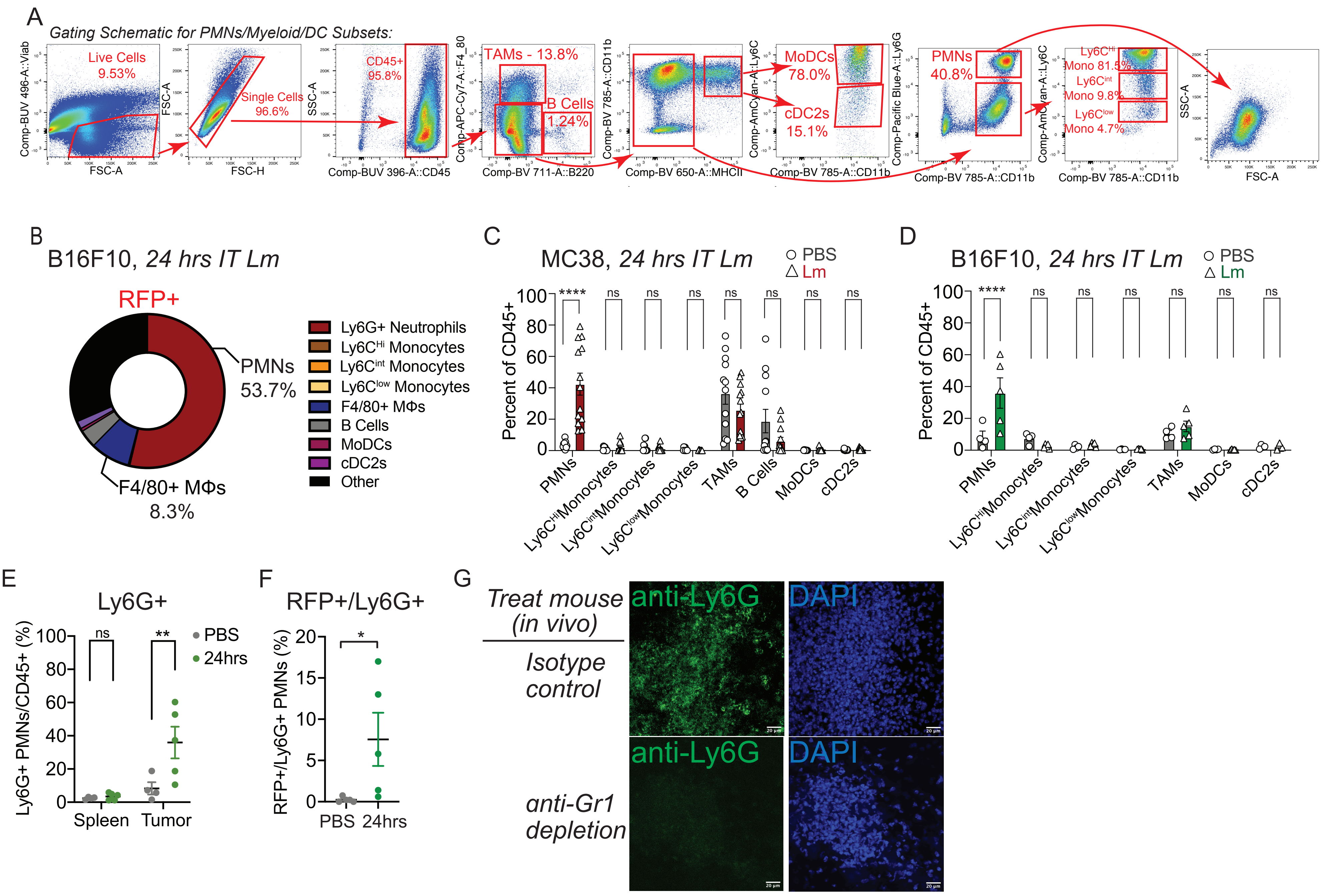


Garia Castillo, J., Campbell, T., & Fernandez, S. et al. Figure S2.

**Figure S2 | Population analysis during intratumoral *Lm* infection in MC38 and B16F10 tumors.**
(A) Gating strategy for identifying immune cell subsets from single cell suspension of MC38 tumor.
(B) Proportion of immune cell types that were $CD45^{+}RFP^{+}$ from B16F10 tumors 24 hours after IT *Lm* (n= 4-5 mice/ group).
(C-D) Frequency of intratumoral immune cell populations as percent of all $CD45^{+}$ cells 24 hours after IT PBS or *Lm* injection in MC38 (C) or B16F10 (D) tumors (n=4-5 mice/group).
(E) Frequency of $Ly6G^{+}$ PMNs in spleen and B16F10 tumors 24 hours after IT PBS or IT *Lm* (n=4-5 mice/group).
(F) Frequency of intratumoral $Ly6G^{+}$ PMNs that were $RFP^{+}$ 24 hours after IT *Lm* in B16F10 tumors.
(G) Validation of anti-Ly6G staining on frozen tumor sections by testing sections from mice treated with an isotype control antibody or an anti-Gr1 antibody that depletes all Ly6G+ cells.
For all plots, mean ± s.e.m. and *P<0.05, **P<0.01, ***P<0.001 by student t-test (B, C, E, F, G, H).

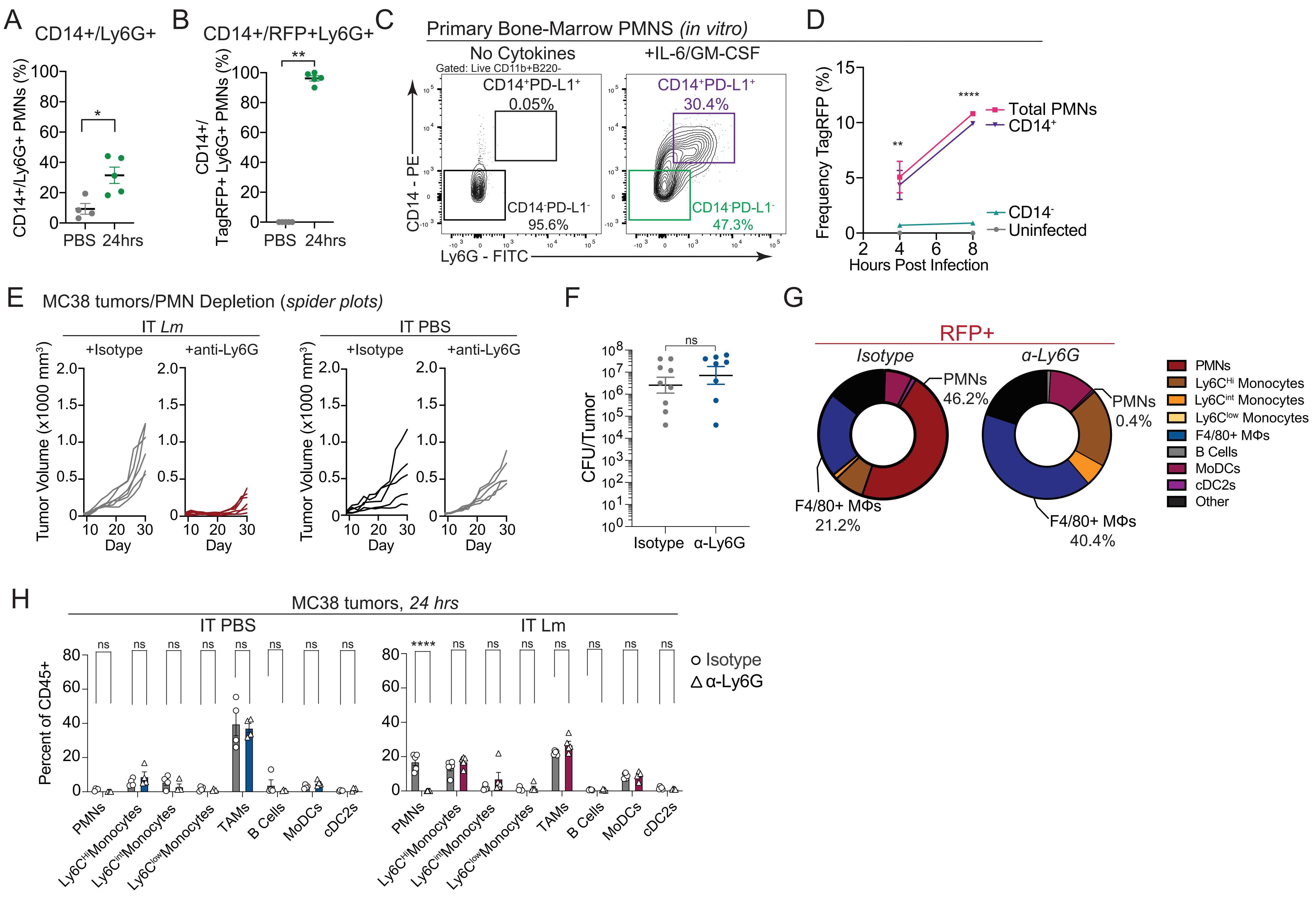


Garcia Castillo, J., Campbell, T., & Fernandez, F. et al. Figure S3.

**Figure S3 | PMN-MDSCs are permissive to *Lm* infection.**

(A) Frequency of CD14+ PMNs 24 hours after IT *Lm* in B16F10 tumors. Results from n= 4-5 mice/group

(B) Frequency of intratumoral CD14+Ly6G$^+$ cells that were RFP$^+$ 24 hours after IT PBS or IT *Lm*.

(C) Representative flow cytometry plots of the differentiation of primary bone marrow progenitors into CD14$^+$ PMN-MDSCs *in vitro* by culturing with IL-6 and GM-CSF.

(D) Heterogeneous PMN-MDSCs differentiated from bone marrow progenitors as in (A) were infected with *Lm-RFP in vitro* and Tag-RFP$^+$ cells were identified 4 or 8 hours later in all cells (Total PMNs), CD14$^+$ cells, or CD14$^-$ cells by flow cytometry.

(E) Tumor growth curves for individual mice in 3J.

(F) *Lm* CFUs recovered from tumors 20 days after IT *Lm* in mice treated with isotype control or anti-Ly6G antibodies as shown in Figure 3J.

(G) Proportion of immune cell types that were RFP$^+$ from MC38 tumors treated with an isotype control (left) or anti-Ly6G antibody (right) 24 hours after IT *Lm*.

(H) Frequency of intratumoral immune cell populations as percent of all CD45$^+$ cells 24 hours after IT *Lm* or IT PBS into MC38 tumors in mice pre-treated with isotype or anti-Ly6G (PMN-depleting) antibodies as shown in Figure 3I.

For all plots, *P<0.05, **P<0.01, ***P<0.001 by student t-test (A-B, F, H), or one-way ANOVA (D), mean ± s.d for *in vitro* experiments. Two-way ANOVA (J), mean ± s.e.m for *in vivo* experiments

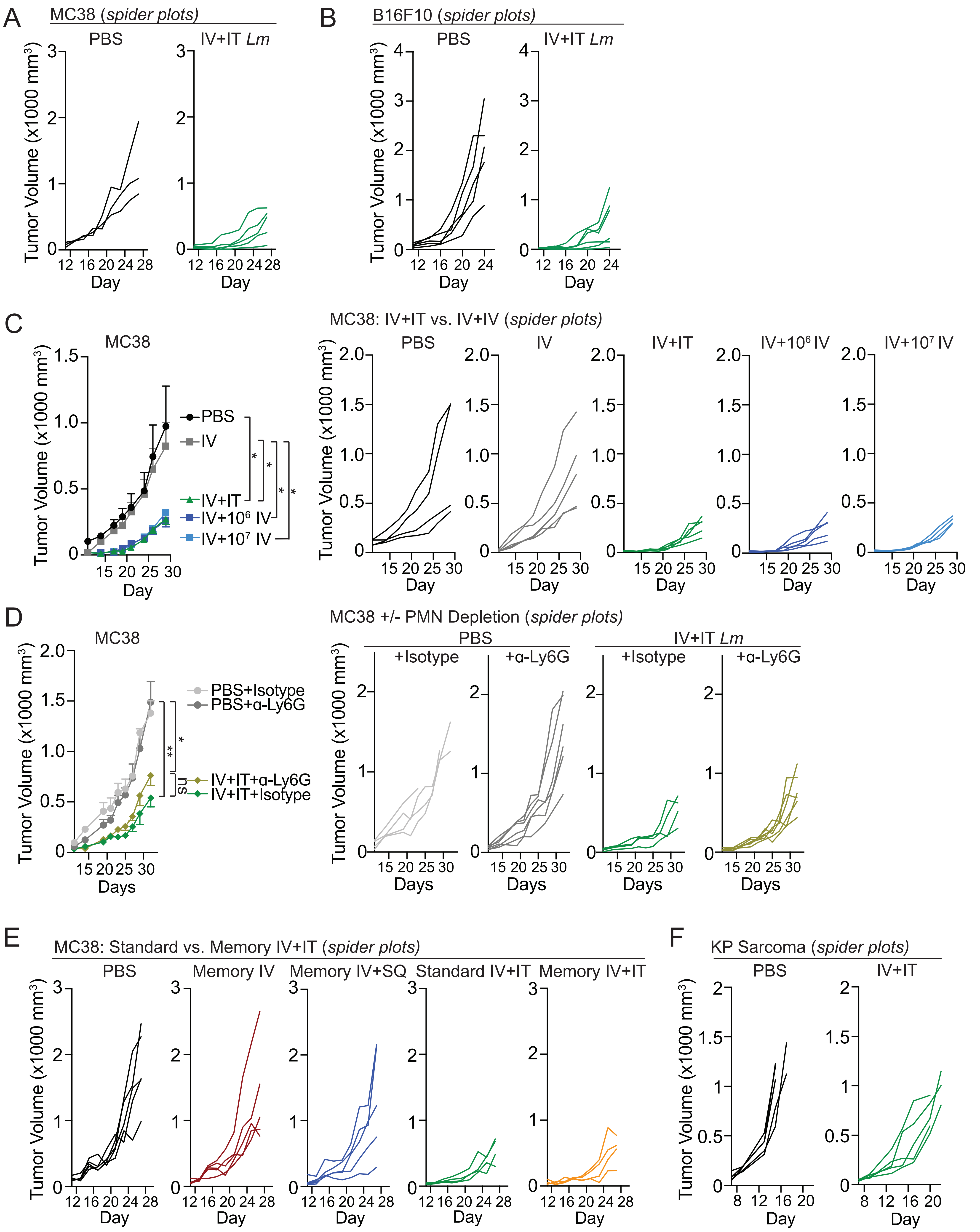


Garcia Castillo, J., Campbell, T., & Fernandez, S. et al. Figure S4.

**Figure S4 | IV+IT *Lm* is not impacted by PMN depletion and IV+IT regimen leads to *Lm* clearance.**
(A) Tumor growth curves for individual mice in 4B.
(B) Tumor growth curves for individual mice in 4C.
(C) Average (right) and individual tumor growth (left) for MC38 with IV+IV PBS versus IV+IV *Lm* versus IV+IT *Lm* regimen (data shown from n= 5-6 mice per group from one of two experiments).
(D) Average (right) and individual tumor growth (left) for MC38 tumor growth with PBS versus IV+IT *Lm* and with or without anti-Ly6G depleting antibody treatment beginning two days prior to IT *Lm* administration (n=4-6 mice per group).
(E) Tumor growth curves for individual mice in 4H.
(F) Tumor growth curves for individual mice in 4I.
For all plots, **P<0.05, **P<0.01, ***P<0.001 by two-way ANOVA (C-D), mean ± s.e.m.

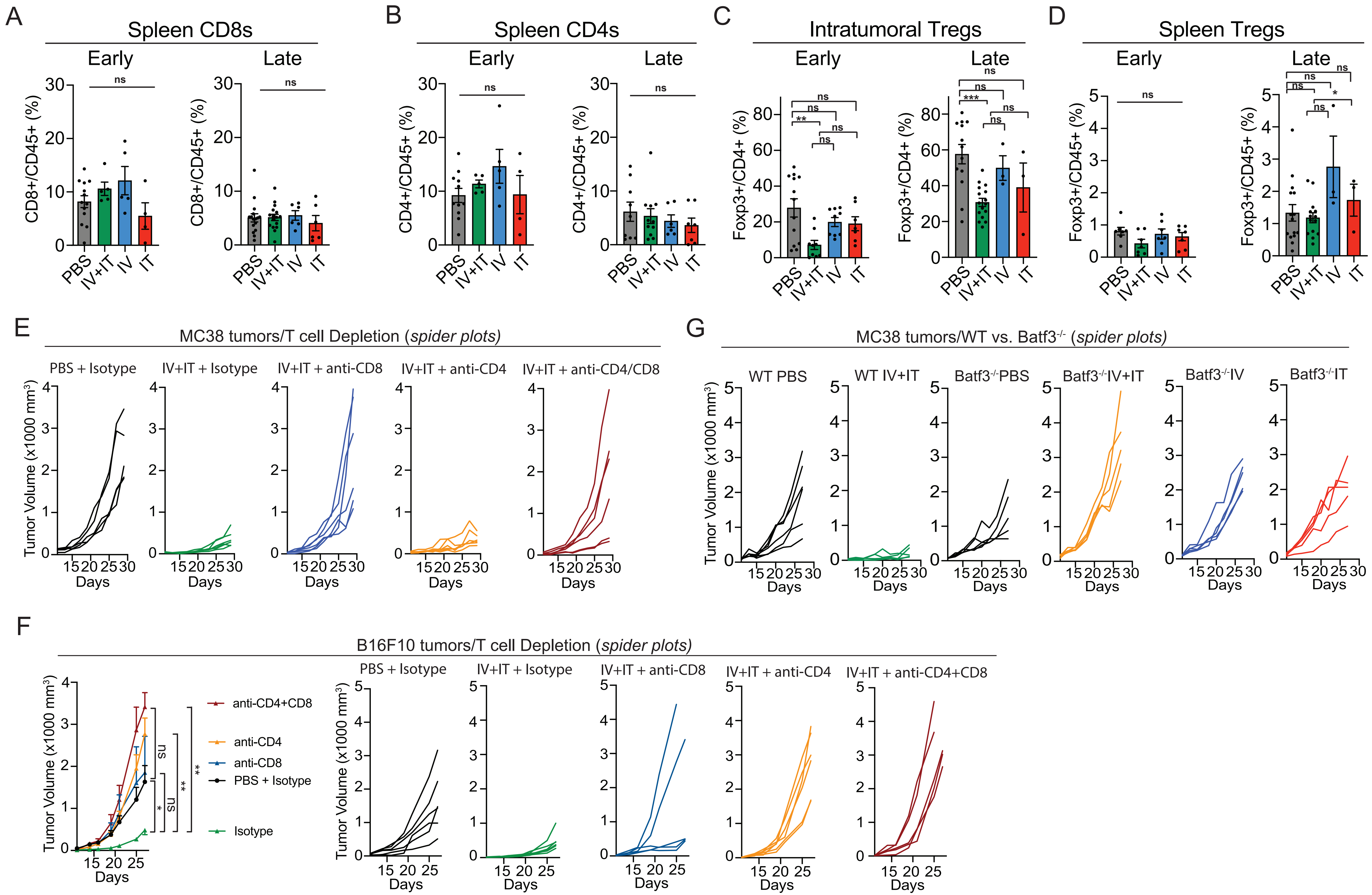


Garcia Castillo, J., Campbell, T., & Fernandez, S. et al. Figure S5.

**Figure S5 | IV+IT *Lm* dosing regimen reduces intratumoral Treg frequencies but does not impact T cell frequencies in the spleen.**

(A-B) Frequency of $CD8^+$ (C) and $CD4^+$ (D) T cells from spleens at early or late timepoints. Data pooled from four experiments with 5 mice/group.

(C) Frequency of intratumoral $Foxp3^+$ Tregs of $CD4^+$ cells from MC38 tumors at early (left) or late (right) timepoints as specified in Figure 5A.

(D) Frequency of $Foxp3^+$ Tregs of $CD4^+$ cells from spleens at early (left) or late (right) timepoints.

(E) Tumor growth curves for individual mice in 5F.

(F) Average (right) and individual tumor growth (left) for B16F10 with IV+IT *Lm* dosing regimen plus anti-CD8, anti-CD4, or both depleting antibodies beginning 10 days after B16F10 tumor inoculation.

(G) Tumor growth curves for individual mice in 5H.

For plots in S5A-S5D, timepoints are the same as what was described in Figure 5A.

For all plots, *P<0.05, **P<0.01, ***P<0.001 by student t-test (A-D) or two-way ANOVA (F), mean ± s.e.m.

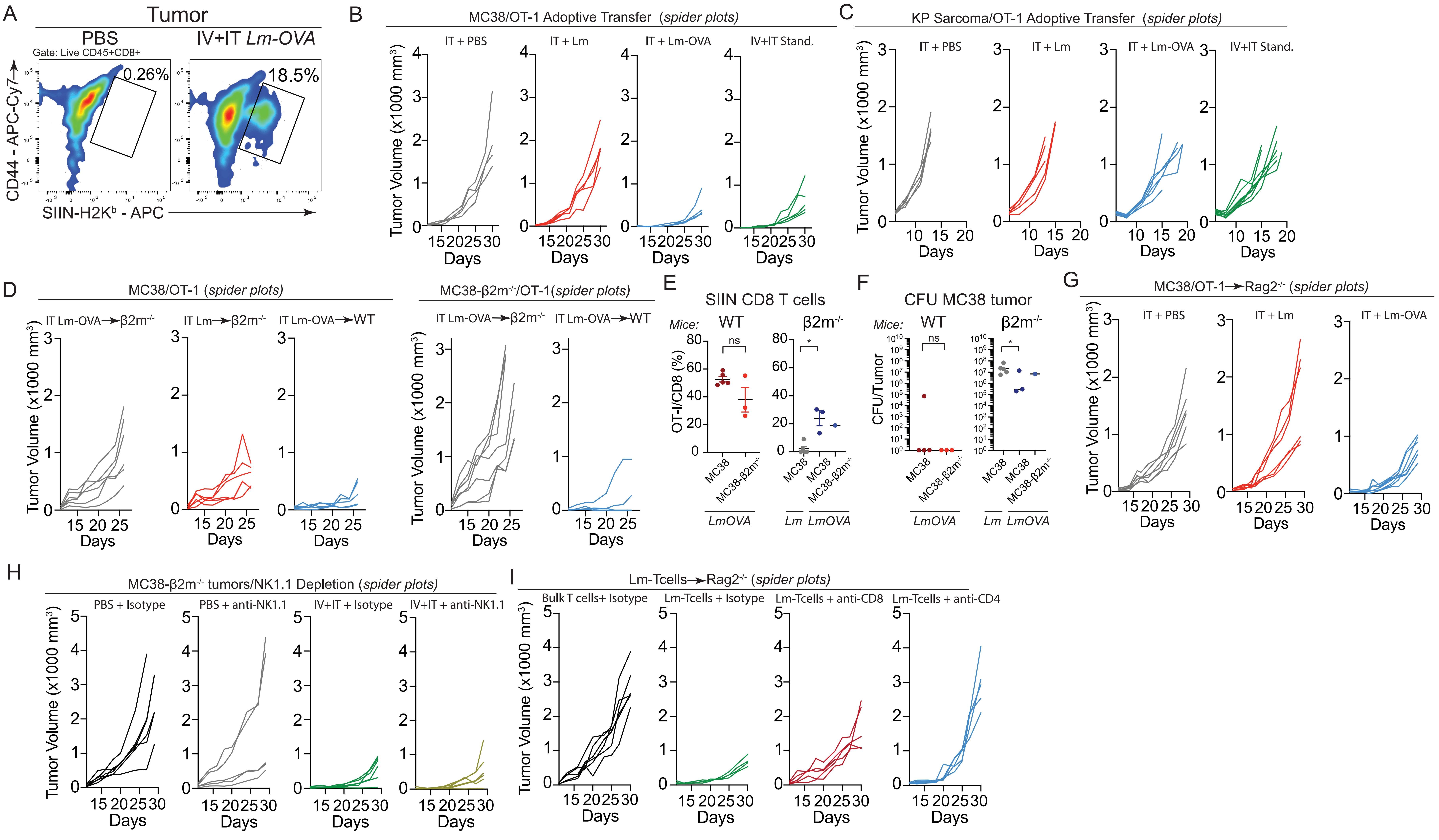


Garcia Castillo, J., Campbell, T., & Fernandez, S. et al. Figure S6.

**Figure S6 | *Lm*-specific CD8 T cells are present in tumors independent of Prf1 genotype.**
(A) Representative flow plots for SIIN-H2K$^b$ tetramer staining from tumors of mice treated with IV+IT PBS compared to IV+IT *Lm-OVA* 8 days after IT injections.
(B) Tumor growth curves for individual mice in 6B.
(C) Tumor growth curves for individual mice in 6C.
(D) Tumor growth curves for individual mice in 6D.
(E) Frequency of intratumoral OT-I CD8 T cells in mice 20 days after IT *Lm* administration from Figure 6D.
(F) CFU analysis for *Lm* in tumors from Figure 6D.
(G) Tumor growth curves for individual mice in 6E.
(H) Tumor growth curves for individual mice in 6F.
(I) Tumor growth curves for individual mice in 6G.
For all plots, *P<0.05, **P<0.01, ***P<0.001 by student t-test (E-F), mean ± s.e.m.

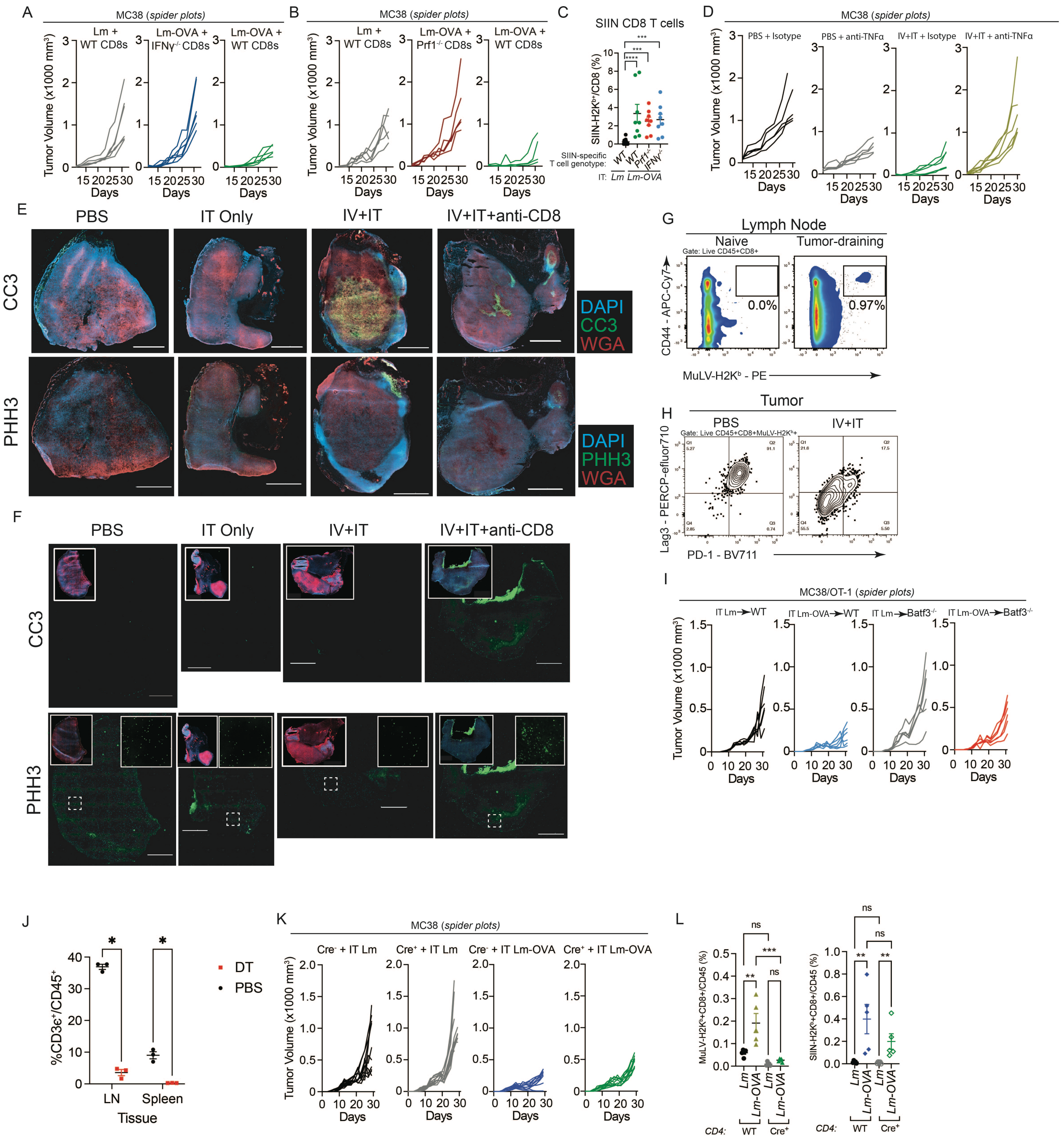

Garcia Castillo, J., Campbell, T., & Fernandez, S. et al. Figure S7.

**Figure S7 | *Antitumor T cell immunity enhancement is dependent on Listeria-specific direct killing and cytokine secretion*.**
(A-B) Tumor growth curves for individual mice in 7B.
(C) Frequency of intratumoral SIIN-specific CD8 T cells in mice 20 days after IT *Lm* administration from experiments depicted in Figures 6J-K (pooled from two independent experiments)
(D) Tumor growth curves for individual mice in 7C.
(E) Images of MC38 tumors 24hrs post injection with IT PBS, IT *Lm*, or given IV+IT dosing regimen +/- anti-CD8 i.p: phospho-Histone 3 (PHH3) or cleaved-caspase 3 (CC3) (green), WGA (red) and DAPI (blue). Scale bar = 2000μm.
(F) Images of MC38 tumors 10 days post injection with IT PBS, IT *Lm*, or given IV+IT dosing regimen +/- anti-CD8 i.p: phospho-Histone 3 (PHH3) or cleaved-caspase 3 (CC3) (green), WGA (red) and DAPI (blue). Scale bar = 2000μm.
(G) Representative flow plots for MuLV-H2K$^b$ tetramer staining in lymph nodes from naive mice compared to MC38 tumor-bearing mice 19 days post tumor inoculation.
(H) Representative flow plots for PD-1+Lag3+ MuLV-H2K$^b$ CD8 T cells for 7G.
(I) Tumor growth curves for individual mice in 7H.
(J) Depletion of T cells in T cell depleter mice (CD4$^{Cre}$; R26$^{LSL-DTR}$) in non-tumor bearing mice. Mice were injected twice with 1μg DT i.p. every two days and inguinal lymph nodes and spleen collected for flow analysis.
(K) Tumor growth curves for individual mice in 7K.
(L) Frequency of intratumoral tumor-specific (MuLV-H2K$^b$ tetramer) CD8 T cells (right) and OT-I CD8 T (left) cells in mice 20 days after IT *Lm* administration from Figure 7J.
For all plots, *P<0.05, **P<0.01, ***P<0.001 by student t-test (C,M) or two-way ANOVA (H), mean ± s.e.m.
